# Trustworthy in silico labeling via semantic visual interpretability of image-to-image translation

**DOI:** 10.64898/2026.08.13.744455

**Authors:** Lion Ben Nedava, Gad Miller, Nitsan Elmalam, Matheus P. Viana, Jianxu Chen, Nathalie Gaudreault, Susanne M. Rafelski, Assaf Zaritsky

**Author notes:** Equal contribution (Lion Ben Nedava, Gad Miller).

## Abstract

Cross-modality image translation promises to provide multiple layers of biological information from a single image input, yet its practical application is stalled by a lack of interpretability and the inability to account for model imperfections. In silico labeling, the inference of organelle localization from label-free images, is a primary example where this black-box nature limits adoption. We present Mask Interpreter, a generalized method for semantic visual interpretability of image-to-image translation models. By uncovering organelle-specific “explanation signatures”, Mask Interpreter validates that models rely on authentic biological structures rather than spurious artifacts. Beyond biological validation, it outperforms traditional explainable AI (xAI) approaches, identifies batch effects and localizes prediction errors when ground-truth fluorescence is unavailable. Semantic confidence modeling further provides fine-grained reliability assessment at single-cell resolution, enabling the automated exclusion of artifacts from downstream analyses. By bridging the gap between computational inference and meaningful biological features, Mask Interpreter transforms in silico labeling into a reliable tool for scientific discovery across diverse biomedical imaging modalities.

## Introduction

Cellular functions involve dynamic and coordinated rearrangements of subcellular organelles and cytoskeletal elements that define the cell’s architecture^1,2^. Thus, uncovering the complex mechanisms of cellular processes requires continuous measurements of a cell’s subcellular organization over time. However, the number of spectrally distinct fluorophores that can be imaged simultaneously in a live cell remains severely limited^3^.

Furthermore, prolonged exposure times, greater imaging depths, or high laser intensities cause photobleaching and induce phototoxicity, altering natural cell physiology^4–6^. Even the attachment of a fluorescent label can disrupt protein structure, alter interactions with binding partners, or impede molecular kinetics, thereby interfering with natural function. Since the 1953 Nobel Prize to Frits Zernike, many label-free imaging methods have been developed for visualizing and measuring subcellular structures from their intrinsic optical properties. In 2018, two seminal studies demonstrated the computational translation of label-free transmitted light (bright-field) microscopy images to synthetic organelle-specific fluorescent images^7,8^, a method coined as ’in silico labeling’ or ‘virtual staining’^9^. These in silico labeling proof of concept studies have been replicated, and applied to new organelles, cell types, and imaging techniques^10–21^. In principle, in silico labeling has the potential to revolutionize cell biology by minimizing the need for fluorescent tags and reducing light exposure. Inferring multiple organelles computationally from label-free images enables longer-term, rapid monitoring of live cellular processes with substantially reduced photobleaching and phototoxicity^9^. However, the real-world application of in silico labeling is currently hindered by several limitations, most notably the risks associated with erroneous localizations, omitted organelles, and model hallucinations^9,19,22^. Thus, its routine deployment remains limited to date to experimental setups where models were optimized to highly specific conditions^20^ or to workflows that can tolerate errors, such as nuclei segmentation^23^ and high-content image-based screening^12,24,25^.

Despite the transformative potential of in silico labeling, a fundamental barrier to widespread adoption remains: the "black box" nature of deep neural networks^9,26^. High performance metrics do not guarantee that a model has learned the underlying biological relationship between label-free structures and organelle localization^27,28^. Much like the classic "husky vs. wolf" dilemma, where a model appeared accurate only because it learned to associate snow in the background with wolves, in silico labeling models may rely on non-biological shortcuts, such as imaging artifacts or batch effects, rather than true subcellular signals^29,30^. This vulnerability is particularly acute when models encounter out-of-distribution data, where the biological state at inference differs from the conditions seen during training. Biological processes such as mitosis^31,32^, differentiation^19^, viral infection^19^, or cell death^33,34^, as well as transitions across cell types or responses to external perturbations (e.g., drug treatments)^19,35–37^, fundamentally alter a cell’s intracellular organization. Even subtler perturbations, such as variations in local cell density or experimental variability (batch effects), can reconfigure the spatial organization of organelles. These shifts change the underlying signal patterns captured in label-free images, which can break the learned transformation and lead to erroneous localizations or hallucinations^9,19,22^. Furthermore, because ground-truth fluorescence is typically unavailable during practical inference, researchers lack a reliable way to verify these predictions or identify where the model’s logic has decoupled from the underlying biology. Without qualitative and quantitative interpretability to distinguish biologically grounded predictions from model failure, in silico labeling cannot yet be trusted for the systematic study of complex cellular processes.

To bridge this gap, we present Mask Interpreter, a generalized method for the semantic visual interpretation of image-to-image translation models. Mask Interpreter uncovers stereotypical explanation signatures within label-free images that drive organelle-specific predictions, revealing the internal logic that deep learning models use to map label-free optical signals to specific subcellular structures. By characterizing these signatures, we demonstrate that identifying anomalous explanation signatures provides a rigorous mechanism for uncovering hallucinations and batch effects without the need for fluorescence ground truth. Supervised quality assessment at inference provides fine-grained reliability metrics that identify where, when, and how in silico labeling succeeds or fails. Semantic confidence modeling allows for single-cell quality control, enabling the automated detection and exclusion of cells with underperforming in silico labeling. This ensures that downstream biological analyses are grounded exclusively in trustworthy in silico labeled cells, providing the oversight necessary to transform in silico labeling into a reliable tool for uncovering new biological insights in complex and dynamic cellular systems. Mask Interpreter is a generalized method for the semantic interpretation of image-to-image translation models that can be readily adapted to other domains in microscopy and beyond.

## Results

### *Mask Interpreter*, a deep-learning model for visual explanations of image-to-image translation models

We used the WTC-11 human induced pluripotent stem cells (hiPSC) Single-Cell Image Dataset v1, created by the Allen Institute. This dataset comprises 3D spinning-disk confocal images of hiPSC lines genetically edited to tag one of 25 key subcellular structures and organelles^38^ (hereafter collectively referred to as organelles). Each 3D fluorescent image had a matching bright-field channel, and a reference nuclear (DNA) and membrane channels. We selected ten organelles with various cell localizations that span the range of in silico labeling performances: nucleoli, nuclear envelope, microtubules, actin filaments, mitochondria, plasma membrane, endoplasmic reticulum, DNA, actomyosin bundles and Golgi (Table S1). We reproduced the in silico labeling results of the corresponding organelles reported in Ounkomol et al.^7^ (Fig. S1) by training organelle-specific U-Net convolutional neural networks^39^ to predict the three-dimensional fluorescence images from the corresponding bright-field image (Fig. 1A).

**Figure 1.**
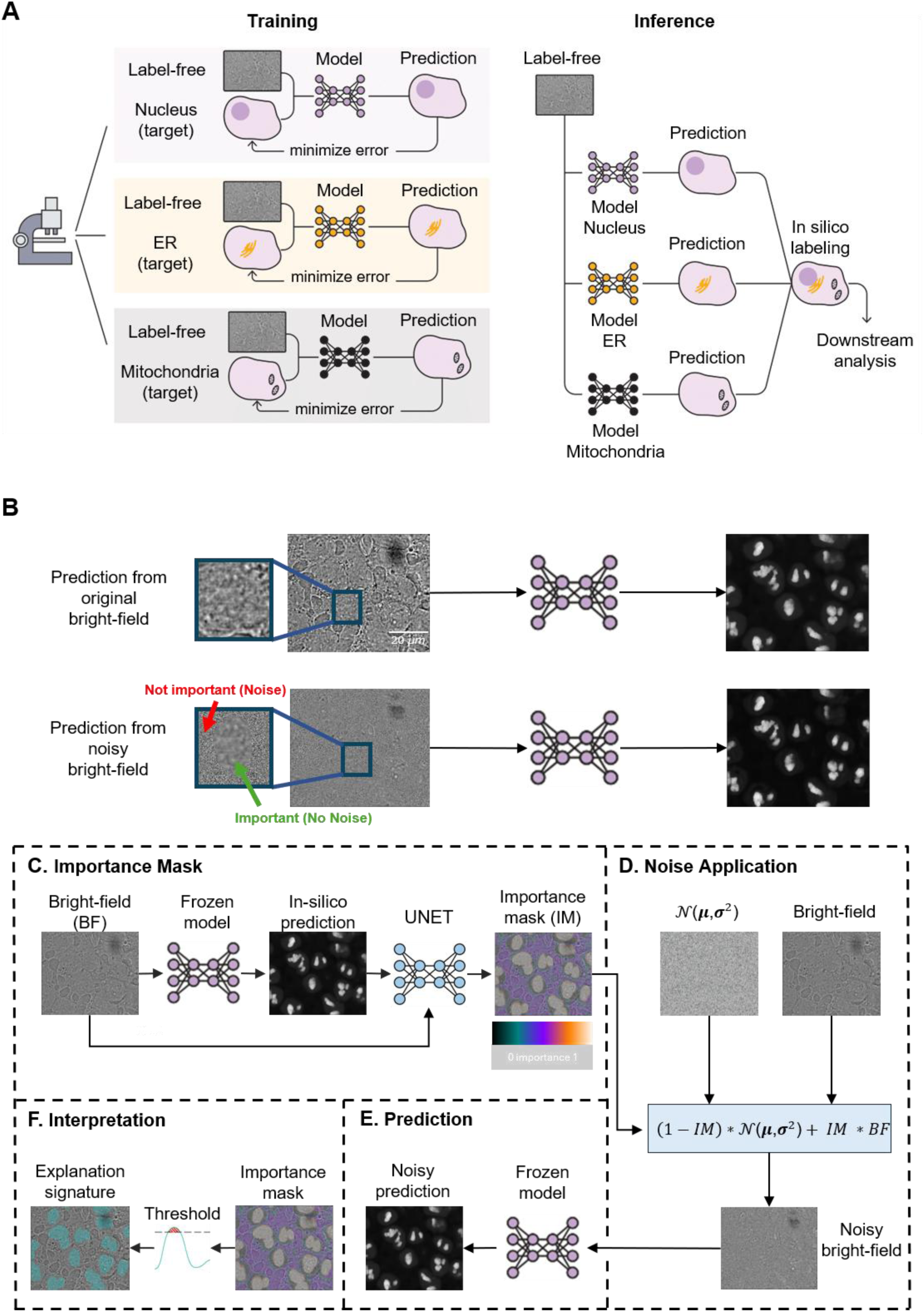
Interpreting in silico labeling using Mask Interpreter. **(A)** In silico labeling. Training (left): organelle-specific in silico labeling models are trained using matched label-free and their corresponding organelle-specific fluorescence images. Inference (right): pre-trained in silico labeling models are used to computationally translate each label-free image to multiple in silico labeled organelles’ fluorescence images. **(B)** An example showing the nucleoli’s in silico labeling from the unperturbed bright-field image (top) and from the noisy bright-field image (bottom). The noisy image is generated by adding Gaussian noise to regions deemed less important by the Mask Interpreter’s derived importance mask. Arrows point to unimportant regions to which noise was added (red) and important regions that were kept intact (green). Note that both in silico labeled predictions are similar, indicating that the in silico labeling model primarily relies on the regions in the bright-free that were deemed as important by Mask Interpreter. Scale bar = 20 μm. **(C-F)** Schematic of the Mask Interpreter training and inference pipeline. (**C**) Importance Mask: The bright-field image is passed through the pre-trained (frozen) in silico labeling model to generate the predicted fluorescence image. The Mask Interpreter then uses the bright-field image and the corresponding prediction to generate a pixel-wise importance mask . Scale bar = 20 μm. (**D**) Noise Application: The noisy bright-field image is constructed by using the importance mask (IM), Gaussian noise (*N*(*μ*, *σ*^2^)), and the original bright-field image (BF) according to (1−IM)×*N*(*μ*, *σ*^2^)+IM×BF. Therefore the noise at each voxel is modulated by the importance mask introducing more noise in less important regions. (**E**) Prediction: The pre-trained in silico labeling model is applied to the noisy bright-field image to generate an in silico labeled prediction. The optimization process minimizes the integrated voxel values in the importance mask (maximizing noise) while simultaneously ensuring the in silico prediction from the noisy bright-field image remains highly correlated (PCC ≥ 0.9) and similar (minimum MSE) to the prediction from the unperturbed bright-field image (see text and Methods). (**F**) Interpretation: The trained Mask Interpreter generates an importance mask from the bright-field. The importance mask is later thresholded to create the binarized explanation signature (see Methods) which provides intuitive visual explanations of the image-to-image translation model’s inner workings.

We developed ‘*Mask Interpreter*’, a general-purpose interpretability method designed to identify the morphological features in a source modality that drive cross-modal translation. We applied this method to reveal the specific visual cues within bright-field images that enable the in silico labeling of an organelle’s location.

Given a bright-field image and its corresponding organelle localization prediction according to a pre-trained in silico labeling model, Mask Interpreter generates an ‘*importance mask*’ explanation image, where voxels that are considered more important for the image-to-image translation are assigned higher values. The determination of a voxel’s “importance” is defined according to the deterioration of the in silico labeling following an alteration in the corresponding voxel in the bright-field image. Voxels with low importance values can be altered by adding noise to the bright-field images, without hampering the in silico labeling performance, while voxels of high importance must remain intact to avoid changes in the in silico labeling prediction (Fig. 1B).

For training, Mask Interpreter requires a pre-trained in silico labeling model that remains frozen throughout the training and a dataset of bright-field images. Mask Interpreter is trained to optimize the weights of an image-to-image translation U-NET architecture that maps a bright-field image and its corresponding in silico prediction to an importance mask (Fig. 1C). The voxels’ values in the importance mask estimate their contribution to the in silico prediction, and during training determine the amount of noise added to the bright-field image. The optimization involves minimizing both the accumulated voxels’ values in the importance mask along with the change in the in silico labeling prediction of the frozen model applied to the noisy bright-field image. More specifically, for a given bright-field image and a given importance mask, Mask Interpreter generates a noisy bright-field image by introducing Gaussian noise at each voxel according to its corresponding value in the importance mask. Voxels with higher importance values are introduced with less noise than voxels with lower importance values creating voxels with higher and lower signal-to-noise ratios correspondingly (Fig. 1D). The noisy bright-field image is in silico labeled using the frozen model (Fig. 1E).

The training relies on backpropagation optimization of three loss terms. The first two complementary loss terms ensure high-quality in silico prediction from the noise-perturbed bright-field image, and were inspired by Liu et al^19^. Specifically, the first loss term is the “target score” loss, enforcing structural fidelity of the in silico labeling from the noisy bright-field image. The target score loss sets the acceptable alteration of the in silico prediction of the noisy bright-field image with respect to the in silico prediction of the unperturbed bright-field image by enforcing a minimum Pearson correlation coefficient (PCC) of 0.9 between them (Methods). The second loss term, “similarity maximization”, enforces intensity consistency between in silico predictions derived from noisy and unperturbed bright-field images by minimizing the mean squared error (MSE) between them. The third loss term is the “mask minimization” that minimizes the integrated voxel values of the importance mask, thereby maximizing the amount of noise introduced into the bright-field image. Together, the target score loss and the similarity maximization loss ensure preservation of both structural features and intensity levels, confirming that the noise introduction does not substantially degrade the in silico labeling. Ablation experiments verified that all loss terms were necessary toward high quality reconstruction (Fig. S2).

Upon inference, the trained Mask Interpreter model computes the importance mask as a quantitative measure of the relative importance of each voxel in the bright-field image to the final in silico labeling prediction. The importance mask can be automatically binary thresholded to highlight the bright-field image regions that are the most important for the in silico prediction (Methods), we call this binary image the ‘*explanation signature*’ (Fig. 1F). These explanation signatures provide human interpretable visual cues of the image-regions that are essential for the in silico prediction^26^.

### Visual interpretation of in silico organelle localization

To interpret the in silico labeling decision process, we applied Mask Interpreter to identify which image-regions within the bright-field input were essential for predicting each specific organelle’s localization. For each of the ten organelle-specific in silico labeling models, we trained a separate instance of Mask Interpreter. Importantly, the hyperparameters for training were calculated using the same pipeline for all ten organelles (Methods). Next, we applied the Mask Interpreter models and visualized their explanation signatures (Methods), to reveal that each organelle had a consistent explanation signature across many cells and fields of view, and that different organelles had different explanation signatures, as described below.

The explanation signature for the nucleoli encompassed the region of the organelle itself and its immediate spatial context visualized at the in-focus focal plane (z-axis) (Fig. 2A, Fig. S3A) and in 3D (Fig. S4A). This explanation implied that the in silico model used local textural patterns in- and adjacent- to the nucleoli in the bright-field image to make its prediction. We found that the in silico mitochondria localization was explained by the organelle along with the spatial context of most of the cytoplasm (Fig. 2B, Fig. S3B, Fig. S4B). The in silico plasma membrane localization was explained by negative spatial context within large image regions that did not include the organelle (Fig. 2C, Fig. S3C, Fig. S4C). Similar dependence on negative spatial context was observed for the actin filaments (Fig. S3D, Fig. S4D) and for the microtubules (Fig. S3E, Fig. S4E).

**Figure 2.**
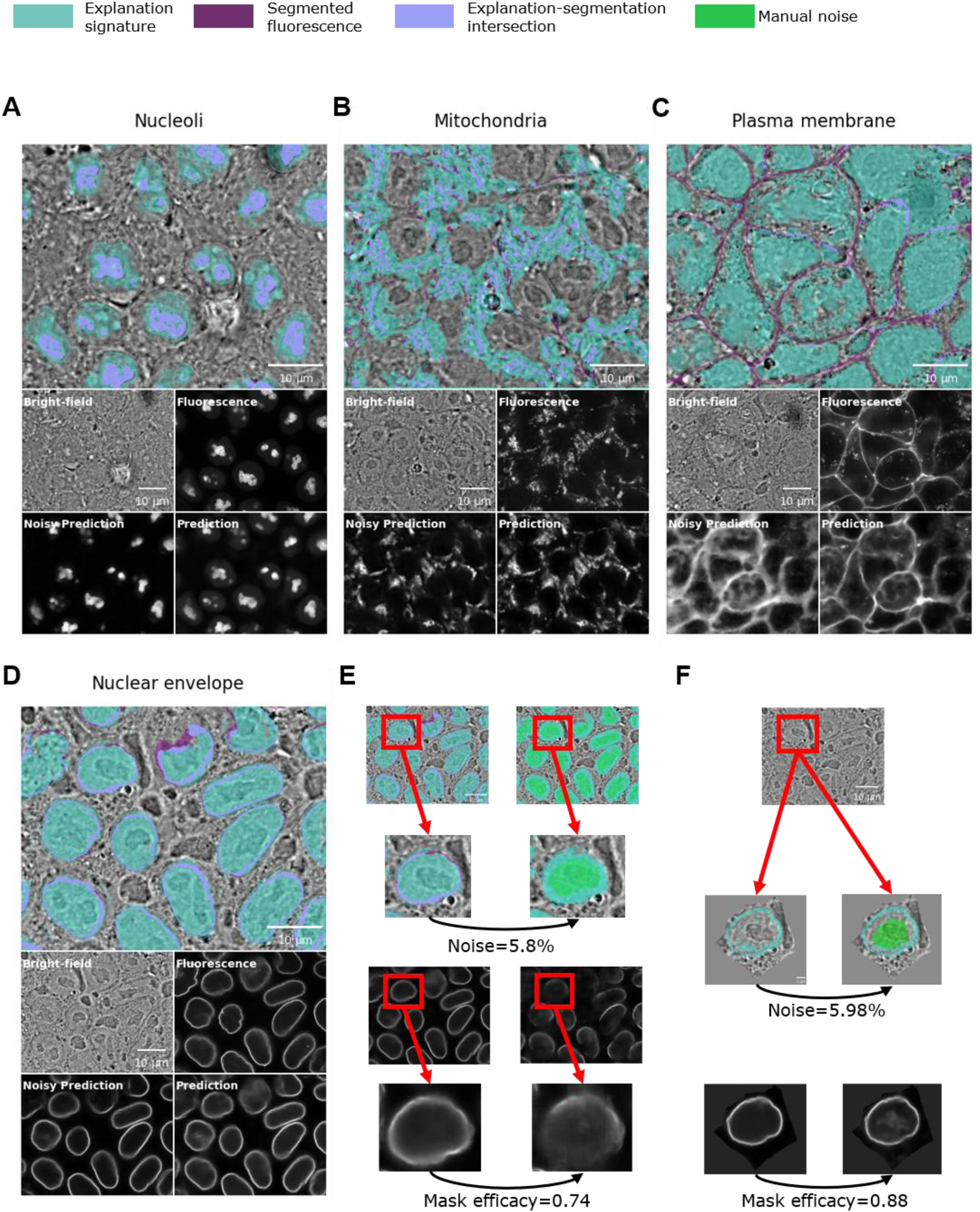
Explanation signatures reveal organelle-specific stereotypical regions that are essential for in silico localization. (**A-D**) A manually selected in-focus z-slice of a representative field of view is shown for each organelle with an overlay of the explanation signature (light teal) and the segmented fluorescence image (purple, Methods). Light purple regions indicate the intersection between the explanation signature and the segmentation (top). The bottom quartet shows the corresponding (top-left to bottom-right) bright-field image, fluorescence image, in silico prediction from the noisy bright-field image (“noisy prediction”) and the in silico labeling from the unperturbed bright-field image (“prediction”). Additional explanation signatures are shown in Fig. S3 (in-focus z-slice) and in Fig. S4 (3D). Explanation signatures of the remaining organelles are shown in Fig. S3 and Fig. S6 (in-focus z-slice) and in Fig. S4 and Fig. S7 (3D). Scale bars = 10 μm. (**A**) The nucleoli. The threshold for the binarization of the importance mask was set to 0.6 (Methods). (**B**) The mitochondria. Binarization threshold=0.2. (**C**) The plasma membrane. Binarization threshold=0.4. (**D**) The nuclear envelope. Binarization threshold = 0.2, leading to a mean noise volume of 81.33% from the field of view measured as the ratio between the number of voxels in the explanation signature and the number of voxels in the image. (**E**) Manual introduction of noise in the bright-field image regions corresponding to the nuclear interior (top, in green) diminished the efficacy of the in silico nuclear envelope prediction, where the mask efficacy was measured by Pearson correlation between the predictions from the noisy and unperturbed bright-filed images (bottom). This direct manipulation underscores the importance of the nucleus in providing the spatial context necessary for accurately localizing the nuclear envelope. Top images scale bar = 10 μm. (**F**) Top: manual introduction of noise in the bright-field image regions corresponding to the nuclear interior that did not intersect with the single cell-centric Mask Interpreter’s explanation signature (cyan). Bottom: the noisy in silico labeling prediction was performed using the single cell-centric in silico labeling model, with minimal effect on the mask efficacy. Additional examples of noise introduction into the nuclear regions in the context of the single cell-centric model are displayed in Fig. S5. Top image scale bar = 10 μm.Other images scale bar = 2 μm.

To further examine how Mask Interpreter resolves distinct spatial dependencies, we performed a detailed analysis of nuclear envelope localization. The predicted nuclear envelope signal was explained by the entire region covered by the nucleus itself, extending to a small margin outside the nuclear boundary (Fig. 2D, Fig. S3F, Fig. S4F). This explanation was surprising to us because the nuclear envelope could be observed as a clear circular gradient in the bright-field image, and thus we expected that the regions associated with the organelle, with some immediate local context around it, would suffice for the in silico labeling. To validate that the region inside of the nuclear envelope that is covered by the nucleus is essential for the in silico prediction, we demonstrated that inclusion of noise in the bright-field image regions where the nucleus resided led to deteriorated in silico labeling (Fig. 2E). The importance of the nuclear region suggested that the nuclear envelope associated patterns in the bright-field image were not nuclear envelope-specific and instead required the nucleus itself to provide spatial context towards accurate in silico localization. We hypothesized that providing the in silico labeling model with a spatial prior of the cells’ center could obviate the need for explicit nucleus-based conditioning for the in silico labeling of the nuclear envelope. To test this hypothesis we trained a single cell-centric in silico model^31^. Briefly, we used the fluorescent plasma membrane-derived segmentation masks to crop single cells, trained an in silico labeling model using these single cell crops and their matched nuclear envelope fluorescence images, and used this model to train a corresponding Mask Interpreter model (Methods). Consistent with our hypothesis, the importance masks derived from this single cell-centric model no longer required the voxels within the nucleus to explain the nuclear envelope localization (Fig. 2F, Fig. S5).

The explanation signatures of the remaining four organelles, the endoplasmic reticulum, the DNA, the actomyosin bundles and the Golgi, appear in Figs. S6-7, and the interpretations of all ten organelles are summarized in Table S2. Overall, these results established that Mask Interpreter can uncover how in silico labeling models transform bright-field images to organelle localization predictions with organelle-specific stereotypical explanation signatures that unveil stereotypical regions in the bright-field image crucial for the model’s predictions.

### Mask Interpreter outperforms alternative visual explanation methods

To validate Mask Interpreter’s explainability capabilities we performed qualitative and quantitative evaluations relative to classic methods for gradient-based “explainable AI” (xAI) that generate heatmaps highlighting the image regions most contributing to a given prediction of deep neural network classifiers. GradCAM uses the gradient information and the activation maps flowing into a specific convolutional layer of the model contributing the most to the output of this layer^40^. Guided backpropagation computes the gradient of the output with respect to the input image^41^. Although both models were designed to explain image-based classification and regression tasks, we applied them to our evaluation due to the lack of established methods for interpreting image-to-image translation. This choice is further justified by viewing image-to-image translation as dense pixel- or voxel-wise regression. The heatmaps generated by these explainability methods can be thresholded and referred to as their corresponding explanation signatures for visualization and for noise introduction in the same manner as in Mask Interpreter. We benchmarked the nucleoli, the nuclear envelope, the mitochondria, and the plasma membrane, because each of these organelles had a different and unique explanation signature. Visualizations of the explanation signatures qualitatively showed that Mask Interpreter generated more intuitive explanations compared to the alternative methods for all four organelles (Fig. 3A-D, top rows). To quantitatively compare the different explanation methods, we evaluated the degradation of the in silico labeling predictions following the inclusion of the same volume of noise in voxels with low importance scores according to the explanation signature derived from each method (Methods). Thus, we devised a new measurement that we term ‘*explanation mask efficacy’* (or in short, ‘mask efficacy’), as the Pearson correlation coefficient between the in silico predictions derived from the unperturbed bright-field images, and the predictions derived from the importance mask-induced noisy bright-field images. A high mask efficacy implies that the in silico predictions of the unperturbed and of the noisy bright-field images are similar to one another because essential voxels were shielded from noise. This analysis suggested that Mask Interpreter explanation signatures encoded the information essential for the in silico prediction better than GradCAM and Guided backpropagation (Fig. 3A-D, middle rows). For example, inclusion of noise in 90% of the lowest-scoring voxels according to Mask Interpreter’s explanation signature of the nucleoli led to a mask efficacy of 0.97 in contrast to inclusion of the same amount of noise, for the exact same image, according to the explanation signatures derived from GradCAM and from Guided backpropagation that dampened the mask efficacy to 0.08 and 0.28, respectively (Fig. 3A). To systematically verify these results we gradually increased the volume of noise according to each explanation method’s importance mask (Mask Interpreter) or heatmap (GradCAM, Guided Backpropagation), and evaluated the mask efficacy between the in silico labeling of the noisy versus the unperturbed bright-field image. This analysis confirmed that the Mask Interpreter consistently tolerated higher levels of noise with substantially less degradation of in silico labeling performance (Fig. 3E). Cumulatively, these results confirm that Mask Interpreter explanation signatures surpass classic xAI approaches in capturing the critical information in the label-free images that was used for the in silico prediction and was thus suitable for gaining insight into the inner decision process of in silico labeling models.

**Figure 3.**
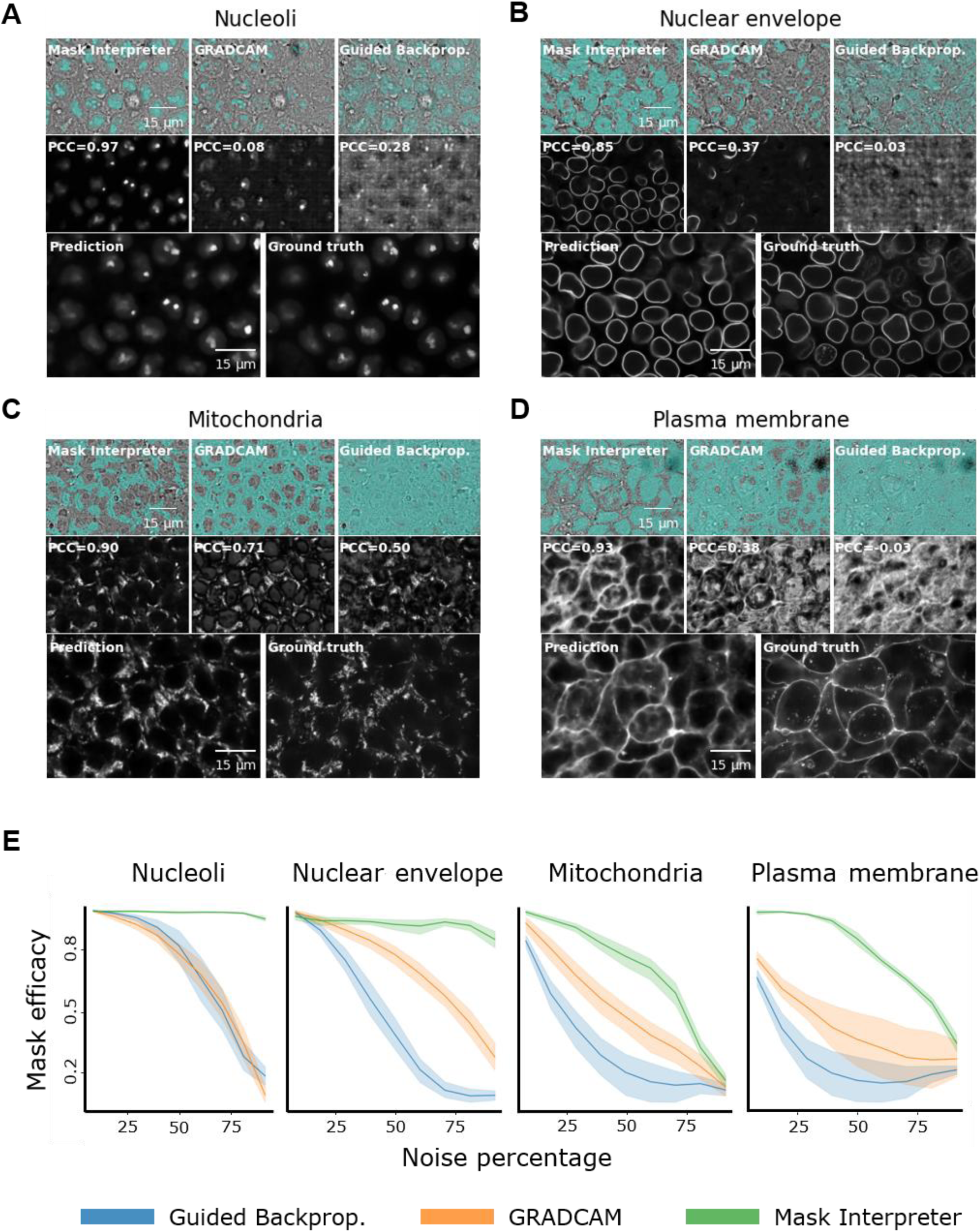
Mask Interpreter surpasses gradient-based interpretability methods for explaining in silico labeling predictions. (**A-D**) Each panel represents a different organelle and includes three rows. Top row: an in-focus z-slice of a representative field of view with an overlay of an explanation signature (cyan) derived from (left-to-right): Mask Interpreter, GradCAM and Guided Backpropagation. The targeted noise percentage was calculated across the complete 3D z-stack rather than independently for each displayed slice; therefore, the visible coverage of the explanation signature may differ between the representative 2D images. This was particularly evident for GradCAM, which frequently assigned high importance to voxels in out-of-focus slices. Middle row: quantitative assessment of the explanation signatures. Comparable amounts of noise were added to the bright-field image according to each method’s thresholded explanation signature and the corresponding in silico labeling of the noisy bright-field image was displayed. The corresponding mask efficacy was reported as a measure of the degradation of the in silico labeling following the introduction of noise, measured by PCC. Bottom row: in silico labeling from the unperturbed bright-field image (left) and the ground truth fluorescence image (right). Scale bars = 15 μm. (**A**) The nucleoli. The explanation signature derived from thresholding the importance mask and used to introduce noise to 90% of the voxels (binarization threshold = 0.6). GradCAM and Guided Backpropagation thresholds were selected to generate explanation signatures with an equivalent 90% noise volume. Corresponding mask efficacies for each of the methods were 0.97 (Mask Interpreter), 0.08 (GradCAM), and 0.28 (Guided Backpropagation). (**B**) The nuclear envelope. Binarization threshold = 0.2 (Mask Interpreter), introducing noise volume of 80%, leading to efficacies of 0.85, 0.37, 0.03 (Mask Interpreter, GradCAM, an d Guided Backpropagation, correspondingly). (**C**) The mitochondria. Binarization threshold = 0.2 (Mask Interpreter), introducing noise volume of 30%, leading to efficacies of 0.9, 0.71, 0.5 (Mask Interpreter, GradCAM, and Guided Backpropagation, correspondingly) (**D**) The Plasma Membrane. Binarization threshold = 0.4 (Mask Interpreter), introducing noise volume of 30%, leading to efficacies of 0.93, 0.38, 0.03 (Mask Interpreter, GradCAM, Guided Backpropagation, correspondingly). (**E**) Systematic analysis on a test set per organelle (up to 10 field of views), demonstrating that Mask Interpreter consistently preserves higher mask efficacies, i.e., the pearson correlation between the predictions from the noisy and unperturbed bright-filed images, as more noise is introduced to the bright-field image according to each xAI method’s explanation signature.

### Altered explanation signatures reveal deteriorated in silico labeling of mitotic cells

Out-of-distribution data, i.e., deviations from the bright-field image patterns observed during training, may be caused by deviations in the intracellular organization during cellular processes, upon perturbations or across cell types. Such deviations may lead to in silico labeling errors, including missed or hallucinated organelle localizations. We hypothesized that anomalous explanation signatures may encode information related to deteriorated in silico labeling due to out-of-distribution data. For example, the nuclear envelope stereotypical explanation signature was a full circular region containing the entire nucleus, the nuclear envelope and some context beyond it (Fig. 2D). In some fields of view we observed explanation masks that did not follow this stereotypical pattern (Fig. 4A) and coincided with degraded disassembled-like nuclear envelope predictions (Fig. 4B-D). In this example, the fluorescent labeled DNA channel resembled a bright elongated blob crossing the cell, characteristic of Metaphase, confirming that the corresponding cell was undergoing mitosis (Fig. 4E). These observations align with prior reports documenting degraded in silico predictions in mitotic cells^32^, particularly for the nuclear envelope^13^. Using the single-cell cell-cycle metadata^38^, we systematically confirmed degraded nuclear envelope predictions in mitotic relative to non-mitotic cells (Fig. 4F), consistent with recent findings^31^.

**Figure 4.**
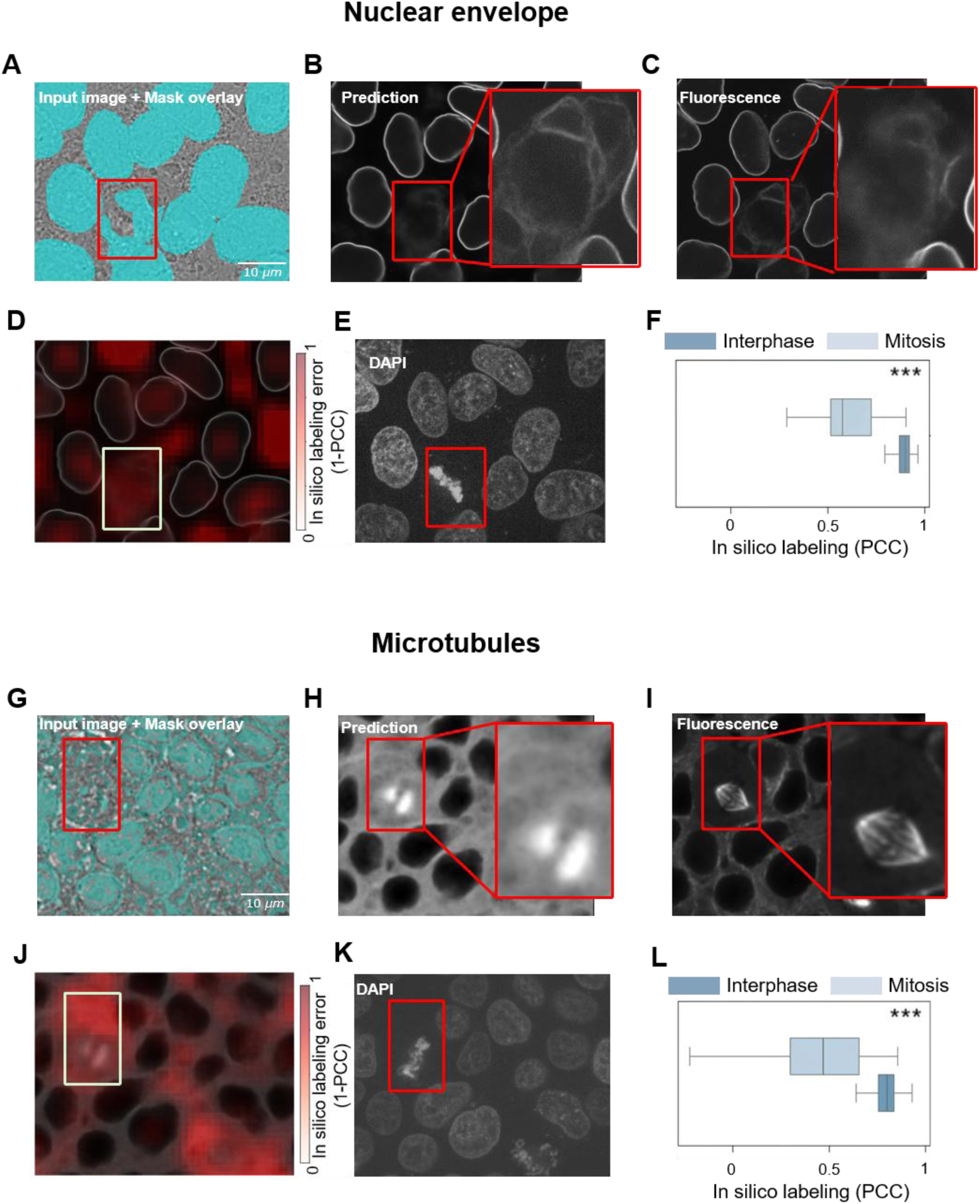
Mask Interpreter reveals non-stereotypical explanation signatures associated with deteriorated in silico predictions for mitotic cells. Non-stereotypical explanation signatures were associated with deteriorated in silico labeling of the nuclear envelope (**A-F**) and of the microtubules (**G**-**L**) in mitotic cells. Scale bar = 10 μm for all panels. (**A-F**) nuclear envelope. Most explanation signatures (cyan) for the nuclear envelope followed a consistent stereotypical pattern, with a minority of cells exhibiting atypical signatures (red bounding box) (A). These atypical cells were associated with diminished in silico labeling accuracy as evident qualitatively and quantitatively between the predicted and ground truth images (B versus C), in the spatial in silico labeling error (1-PCC) correspondingly (D). The DNA fluorescence channel confirmed that the corresponding cell was undergoing mitosis (E). Systematic evaluation across n = 189 cells from N = 16 held-out fields of view confirmed that mitotic cells (light blue) displayed deteriorated in silico nuclear envelope labeling performance compared with interphase cells (dark blue). Box plots show single-cell PCC (Methods): median (center line), first and third quartiles (box bounds), and whiskers extending to minimum and maximum values within the 1.5× the interquartile range. One-sided Mann–Whitney U test, *** p-value < 0.001 (F). **(G–L)** Microtubules. Details in panels G-L correspond to those described in A-F. Non-stereotypical explanation signatures for the microtubules were observed in mitotic cells (G). These atypical signatures corresponded to lower in silico labeling performance qualitatively (H versus I), and quantitatively (J). The DNA fluorescence channel confirmed that the corresponding cell was undergoing mitosis (K). Systematic evaluation across n = 145 cells from N = 11 held-out fields of view confirmed that mitotic cells displayed deteriorated in silico microtubules labeling compared with interphase cells, p-value < 0.001 (L).

This degradation suggests that the low abundance of cells undergoing mitosis in the training datasets limits the model’s ability to generalize to mitotic bright-field morphology. Similarly, microtubules exhibited non-stereotypical explanation signatures coupled with degraded predictions in mitotic cells (Fig. 4G-L)^31^. These results suggested that anomalous explanation signatures could flag deteriorated in silico labeling.

### Mask Interpreter uncovers batch effects in bright-field microscopy

Batch effects are technical variations that can arise from differences in factors such as microscopes, protocols, laboratories, experimentalists, and cell seeding densities^42,43^. In the context of this dataset, because FOVs for each of the genetically edited cell lines were acquired under the same imaging pipeline, we defined each cell line as an individual ’imaging batch’. Systematic variations between these imaging batches can introduce out-of-distribution optical shifts in bright-field images, leading to batch-dependent in silico labeling errors. We speculated that Mask Interpreter could be used to detect such batch effects at inference. Our hypothesis was that deviations from the bright-field image distribution in the training data would lead to explanation masks that fail to encapsulate the essential image regions toward the in silico prediction. If true, deviations from the bright-field image will lead to low mask efficacies (Fig. 5A). Importantly, mask efficacy can be calculated at inference, even when the ground truth fluorescence images are not available. Additionally, the “target score” loss term enforces an explanation mask’s efficacy of ∼0.9 during training, and thus, mask efficacies substantially lower than 0.9 indicate that the in silico labeling model cannot accommodate the added noise according to the explanation mask, suggesting that the bright-field image may be out-of-distribution (Fig. 5B). Thus, reduced mask efficacies should serve as an indicator of batch effects that impair in silico labeling.

**Figure 5.**
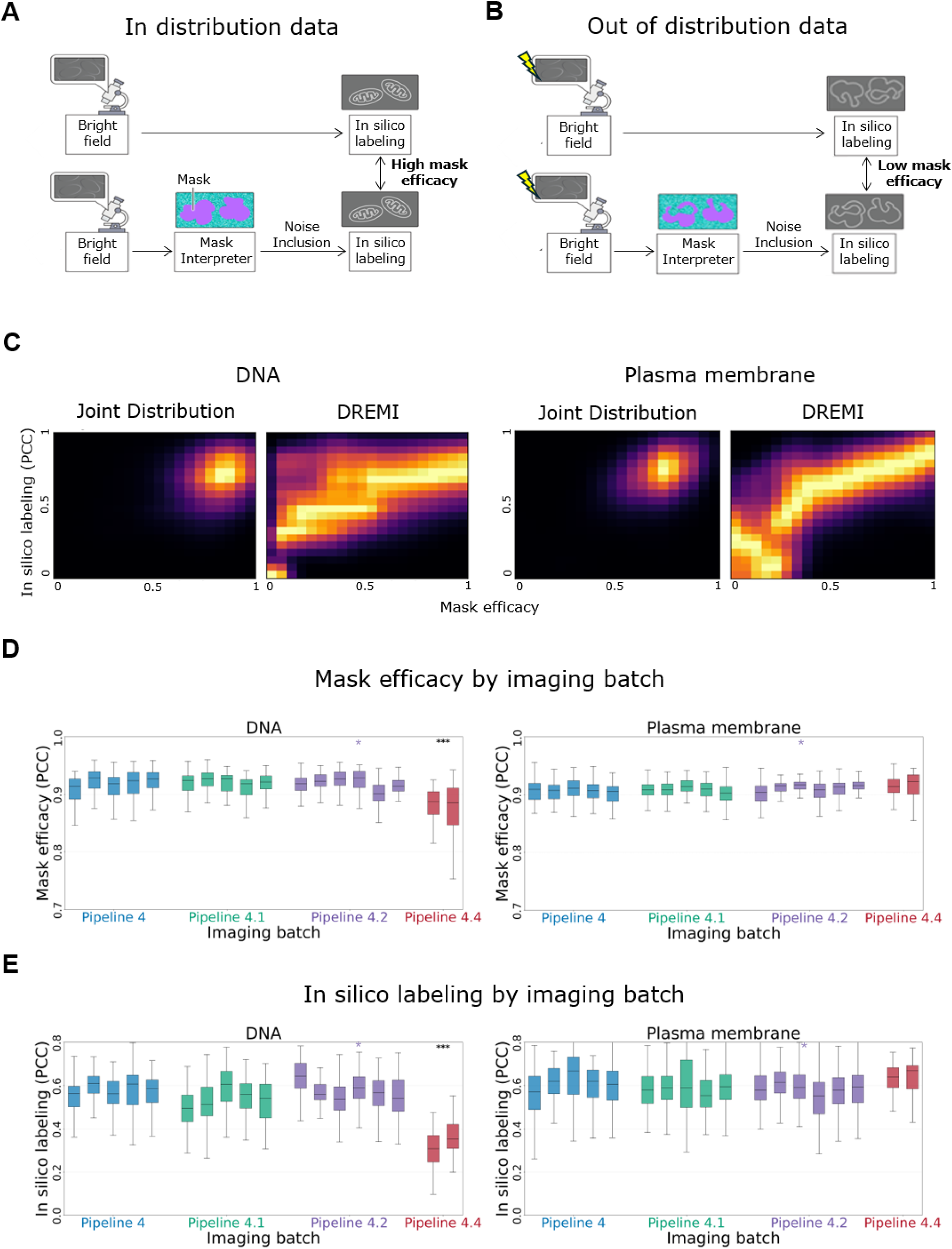
Mask Interpreter detects batch effects that impair in silico labeling performance. **(A, B)** Conceptual schematics illustrating the mask efficacy measurement for in-distribution (A) and out-of-distribution (B) data. (**A**) When bright-field images are in-distribution, introduction of noise according to the explanation mask induces minimal changes in the in silico labeling, leading to a high mask efficacy score. (**B**) In contrast, we hypothesized that out-of-distribution data will lead to explanation masks that fail to encode the bright-field information essential for the in silico labeling. This will lead to deviations between the unperturbed and the noisy in silico labeling predictions, and correspondingly to a low mask efficacy score. **(C)** Joint distribution and conditional density (DREMI) between the mask efficacy (x-axis) and the in silico labeling (y-axis) pooled across N = 1,821 fields of view for the DNA (left) and for the plasma membrane (right). The joint distribution concentrated in a narrow range of values confounding the true relationship between these measurements across their full dynamic range with mutual information = 0.05 and 0.12 for the DNA and for the plasma membrane, correspondingly. Density-corrected DREMI scores (Methods) reveal stronger associations between the mask efficacy and the in silico prediction of 0.26 and 0.53 for the DNA and for the plasma membrane, correspondingly. This result indicated that decreased mask efficacy was associated with degraded in silico labeling. (**D**) Mask efficacy across imaging batches and pipelines. Box plots display FOV-level mask efficacy distributions for DNA (left) and plasma membrane (right). Each box represents a single imaging batch, with colors denoting distinct imaging pipelines (Pipelines 4, 4.1, 4.2, and 4.4). Center lines indicate the median, boxes span the interquartile range, and whiskers extend to 1.5 × the interquartile range (outliers omitted). In silico models and corresponding Mask Interpreter models were trained on a subset of FOVs from a single batch (marked with a purple asterisk) within Pipeline 4.2. Statistical comparisons were performed using a one-sided Mann–Whitney U test, comparing the held-out test FOVs of the training batch against the pooled FOVs across all batches within each respective pipeline. For DNA, the held-out test set included N = 15 FOVs, and the evaluated pipelines comprised N = 641 (Pipeline 4), N = 465 (Pipeline 4.1), N = 446 (Pipeline 4.2), and N = 116 (Pipeline 4.4) total FOVs. The DNA model exhibited a significant drop in mask efficacy specifically for Pipeline 4.4 (p = 7.3 × 10^-7^), flagging a pipeline-specific shift in the bright-field image distribution. **(E)** In silico labeling performance across imaging batches and pipelines. Box plots display FOV-level in silico labeling distributions for DNA (left) and plasma membrane (right). Each box represents a single imaging batch, with colors denoting distinct imaging pipelines (Pipelines 4, 4.1, 4.2, and 4.4). Center lines indicate the median, boxes span the interquartile range, and whiskers extend to 1.5 × the interquartile range (outliers omitted). In silico models and were trained on a subset of FOVs from a single batch (marked with a purple asterisk) within Pipeline 4.2. Statistical comparisons were performed using a one-sided Mann–Whitney U test, comparing the held-out test FOVs of the training batch against the pooled FOVs across all batches within each respective pipeline. For DNA, the held-out test set included N = 15 FOVs, and the evaluated pipelines comprised N = 641 (Pipeline 4), N = 465 (Pipeline 4.1), N = 446 (Pipeline 4.2), and N = 116 (Pipeline 4.4) total FOVs. The DNA model exhibited a marked decreased in silico labeling performance specifically for Pipeline 4.4, corresponding to a 41% performance reduction (p-value = 4.5e-10). Plasma membrane in silico labeling did not show deterioration of the in silico labeling performance. Results for eight additional organelles are presented in Fig. S8 and in Fig. S9.

To systematically assess this hypothesis, we evaluated the variation in mask efficacy and corresponding in silico labeling performances for the reference nuclear (DNA) and membrane channels across the different imaging batches acquired over more than a year, across different microscopes, and through evolving imaging pipelines. The DNA and membrane were selected because they were tagged and imaged in every imaging batch. Mask efficacy scores were computed for the in silico labeling of 1,821 fields-of-view (FOVs) pooled across imaging batches. Direct estimation of the mutual information *P*(*X*, *Y*) between mask efficacy and in silico labeling performance obscures their true dependency across their full dynamic range because their values concentrate in a narrow interval (Fig. 5C - joint distribution). To resolve this, we employed conditional-Density Resampled estimate of Mutual Information (DREMI)^44^, which estimates the conditional density *P*(*Y*|*X*) by distributing FOV mask efficacies into evenly spaced bins along the x-axis, thereby rebalancing dense and sparse regions of the distribution (Methods). DREMI revealed a strong association between mask efficacy and in silico labeling performance (Fig. 5C - DREMI).

Evaluating the different imaging batches revealed that DNA mask efficacies in two imaging batches were strikingly low compared to the rest (Fig. 5D). Metadata analysis traced this discrepancy to a deviation in the image acquisition pipelines. Specifically, the Single-Cell Image Dataset v1 was generated using four distinct imaging workflows that were marked as pipelines 4.0, 4.1, 4.2, and 4.4. While the first three pipelines used a broadband halogen light source and a single-camera setup, pipeline 4.4 employed a far-red LED light source and a dual-camera system. These low mask efficacies observed in the two imaging batches acquired with pipeline 4.4 (Fig. 5D) directly corresponded to a drop in DNA in silico labeling (Fig. 5E). Conversely, plasma membrane mask efficacies remained unaffected under pipeline 4.4, with no corresponding performance deterioration (Fig. 5D-E). These results demonstrate that hardware and acquisition changes can differentially impact the in silico labeling of different organelles. Encouraged by this ability of mask efficacy to capture the differential performance between DNA and plasma membrane predictions, we extended our analysis to the remaining eight organelle models. For five of these models, mask efficacy yielded markedly low scores when applied to bright-field images acquired under pipeline 4.4 (Fig. S8). Notably, because fluorescence ground truth was available only for the specific pipeline used during model training, cross-pipeline predictions lacked paired fluorescence data, meaning these predicted drops in efficacy could not be quantitatively verified and were restricted to visual assessment (Fig. S9).

### Supervised quality assessment at inference

Mask efficacy was shown to identify deteriorated in silico labeling of fields of view in an unsupervised manner. We next reasoned that performance estimation could be improved by incorporating direct supervision, specifically by training a regression model to predict local prediction accuracy. The success of the mask efficacy measurement also led us to speculate that the importance mask can provide relevant information for predicting the model’s performance, along with providing the trustworthiness of the “built-in” explanation signature. Thus, we fine-tuned a 3D ResNet^45^ “confidence” model to predict the in silico labeling performance, as defined by the per image-patch Pearson correlation between the in silico labeling prediction and the corresponding fluorescent image (Methods). Operationally, the model was trained to predict the corresponding prediction error, defined as (1 - this confidence). The confidence model received a two-channel input comprising the in silico labeled patch and its corresponding Mask Interpreter importance mask (Fig. 6A). Training was performed across eight organelles, excluding the actomyosin bundles and Golgi, whose poor in silico labeling performance limited the utility of confidence estimation. The model was trained to predict the in silico labeling performance by minimizing Mean Squared Error. To compare the supervised confidence model against the unsupervised mask efficacy score, we performed a paired patch-level analysis. For each patch, we defined the *estimation error* as the absolute difference between the predicted performance score and the actual performance (measured as the patch-level PCC against fluorescence ground truth). Subtracting the supervised model’s estimation error from that of mask efficacy yielded positive values, demonstrating that the confidence model provided more accurate performance estimates (Fig. 6B; Methods). Evaluating these errors via Mean Squared Error (MSE), PCC, and R^2^ confirmed that the supervised model yielded consistently lower estimation errors, systematically outperforming mask efficacy across all organelles (Tables S3-S5; Methods). DREMI analysis confirmed superiority of the supervised confidence model for six of the eight organelles (Fig. S10). An ablation analysis showed that including the importance mask provided mild additional predictive information beyond the in silico labeling output alone, with improved performance in five of the eight evaluated organelles (Fig. S11). The full results are summarized in Tables S3-S5. Visual assessment of image regions that were predicted as poor in silico labeled further verified the spatial association with the ground truth errors in the in silico labeling, suggesting that the confidence model is capable of identifying in silico labeled regions that underperform without access to ground truth data (Fig. 6C, Fig. S12). Systematic DREMI analysis confirmed the capacity to predict the in silico labeling performance at the patch level during inference, specifically identifying the image patches that cannot be trusted for downstream analysis (Fig. 6D, Fig. S13). Together, these results establish that supervised confidence modeling provides fine-grained reliability assessment at inference, directly complementing Mask Interpreter’s interpretability.

**Figure 6.**
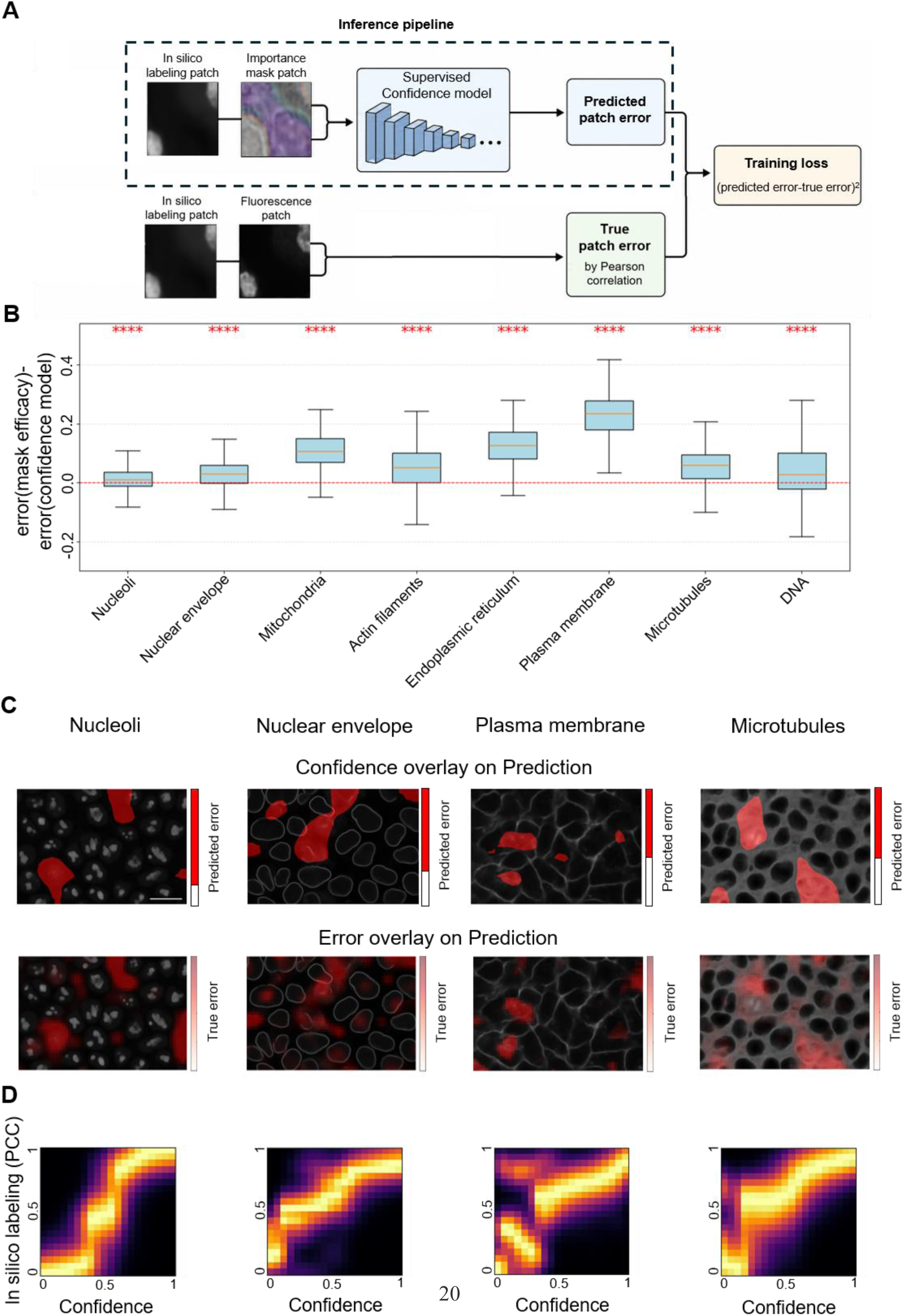
Supervised confidence model predicts unreliable in silico labeled regions. **(A)** Schematic of the confidence model pipeline. The supervised confidence model receives as input the in silico labeling of an image patch and its corresponding Mask Interpreter-derived importance mask. The model is trained to predict the in silico labeling performance. The model is optimized to minimize the mean squared error (MSE) between the predicted and the actual in silico labeling performance, defined as the PCC between the fluorescence and the in silico labeling. **(B)** Comparison between the supervised confidence model and the mask efficacy measurement. For each organelle, performance differences were computed in a paired model-comparison analysis. The y-axis shows the difference in absolute PCC error between the mask efficacy and the confidence model’s estimation errors. Positive values indicate better performance of the supervised confidence model. The statistical test is performed on the patch level (left-to-right, N=810, 540, 972, 540, 378, 432, 324, 810) and evaluates whether the observed performance difference between the compared confidence measures/models is larger than expected by random variation. ****P < 0.0001. Box plots show the median and interquartile range (IQR); whiskers extend to the most extreme values within 1.5 × IQR. The red dashed horizontal line at zero marks that both models had the same estimation errors. The full results are available in Tables S3-S5. **(C)** Visualization of predicted and ground-truth-verified low-quality in silico labeling regions. Representative fields of view are shown for (left-to-right), nucleoli, nuclear envelope, plasma membrane, and microtubules. In the top row, manually-defined threshold (left-to-right: 0.18, 0.3, 0.42, 0.4) low-confidence regions predicted by the supervised confidence model are overlaid in red on the in silico labeled image. In the bottom row, low-quality regions, identified by comparison to the corresponding fluorescence ground truth, are overlaid in red on the same in silico labeled image. Scale bar = 15 μm. (**D**) Systematic evaluation of the confidence model for the (left-to-right) nucleoli (810 patches), Nuclear Envelope (540 patches), Plasma Membrane (432 patches) and microtubules (324 patches) using DREMI analysis of the predicted versus ground truth in silico labeling PCC. DREMI scores (left-to-right): 0.88, 0.62, 0.43, 0.51.

### Semantic confidence at single-cell resolution enables automated quality control

Individual cells are the fundamental unit of many downstream analysis pipelines. We therefore extended our evaluation beyond patch-level predictions to assess in silico labeling performance at the single-cell level. Such single-cell quality control can flag and filter algorithmic artifacts introduced by underperforming predictions^46,47^, ensuring that downstream biological inferences are derived exclusively from trustworthy cells and organelles.

Because general improvements in pixel-level image quality metrics do not always translate to improved performance in downstream analysis^28^, we focused on “application-appropriate” metrics suited for biology-driven quantification^48^, specifically, the downstream task of organelle segmentation^49^. Single-cell segmentation masks provided by the Allen Institute’s dataset were used to isolate individual cells (Fig. 7A, Methods). We focused this analysis on the nucleoli, nuclear envelope and microtubules, as these organelles yielded consistent and biologically meaningful segmentations using the Allen Institute’s Structure Segmenter pipeline^77^. For each organelle, we applied the standard segmentation pipeline identically to both the in silico labeling prediction and their corresponding ground-truth fluorescence images. Paired segmentations were evaluated by calculating the Intersection-over-Union (IoU) and Dice Coefficient across single cells. DREMI analysis between the predicted confidence scores and these segmentation accuracy metrics (IoU, Fig. 7B; Dice coefficient, Fig. S14A) confirmed that lower predicted confidence corresponds to reduced segmentation quality. Stratifying single cells into quartiles according to their predicted confidence scores revealed a pronounced drop in segmentation fidelity within the lowest-confidence quartile (‘Q1’; Fig. 7C-D, Fig. S14B). These results demonstrate that supervised confidence scores provide a practical, single-cell quality-control mechanism to automatically identify and exclude unreliable in silico predictions prior to downstream analysis.

**Figure 7.**
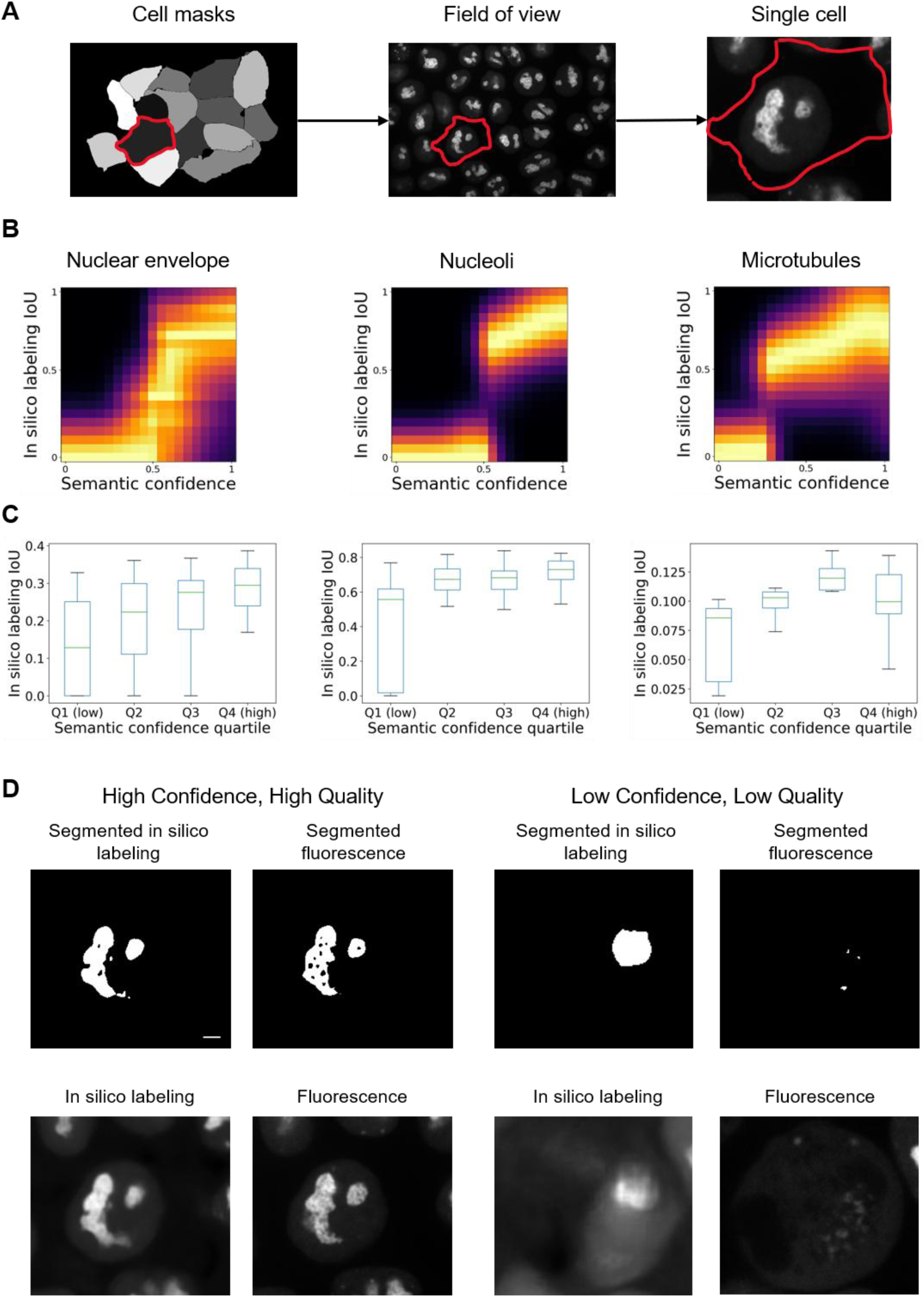
Single cell semantic confidence identifies poor in silico labeling for organelle segmentation-based downstream analysis. **(A)** Schematic of extracting single-cell images. Field of view (FOV) cell masks provided by the Allen Institute dataset^38^ (left) were used to extract in silico labeling of single cells (right) from the FOV (middle). The single cell semantic confidence was calculated by averaging the predicted confidence over the contiguous 16-slice z-window with the highest summed in silico-labeling intensity within the cell mask (Methods). **(B-C)** Single cell semantic confidence associates with intersection over Union (IoU) between the segmented in silico labeling and segmented fluorescence image. The corresponding Dice Coefficient analysis appears in Fig. S12. (**B**) Conditional density (DREMI) between the single cell semantic confidence (x-axis) and the IoU between the segmentations of the in silico labeling and the fluorescence ground truth, across three organelles (left-to-right): Nuclear Envelope (143 cells, DREMI score = 0.45), nucleoli (176 cells, DREMI score = 0.81.), and microtubules (33 cells, DREMI score = 0.51). (**C**) Cells were grouped into quartiles (Q1–Q4) based on their semantic confidence scores, with Q1 corresponding to the lowest confidence and Q4 to the highest. Boxes indicate the interquartile range, center lines indicate the median, and whiskers extend to extreme values within the 1.5× the interquartile range (outliers not shown). IoU was higher in the pooled Q2–Q4 cells than in Q1 cells for all three organelles: Nuclear Envelope (median: 0.274 versus 0.128; one-sided Mann–Whitney U = 2883.5, p = 4.0 × 10⁻⁶; Cliff’s δ = 0.497), nucleoli (0.691 versus 0.557; U = 4970, p = 8.51 × 10⁻¹³; Cliff’s δ = 0.711), and microtubules (0.107 versus 0.086; U = 182, p = 1.48 × 10⁻³; Cliff’ s δ = 0.685; note lower overall IoU values for microtubules reflect their sparse, filamentous structure). **(D)** Representative nucleoli examples from a high semantic confidence cell with strong agreement between the segmented in silico labeling and fluorescence image (left) and a low semantic confidence Q4 cell with poor segmentation agreement (right). Scale bars = 2 μm.

## Discussion

The challenge of establishing "trustworthy AI" has emerged as a fundamental bottleneck in the integration of deep learning into the high-stakes domains of science and medicine. Trust is not merely a measure of predictive accuracy, but a requirement for ensuring that models are right for the right reasons, particularly when deployed in out-of-distribution settings where ground-truth verification is impossible. Without transparency, even high-performing models remain "black boxes" susceptible to hallucinations, batch effects, and the systematic propagation of errors into downstream applications, which can lead to erroneous conclusions or the dismissal of genuine, unexpected phenomena. Interpretability is the primary vehicle for resolving this challenge, providing the necessary oversight to transform AI from an opaque predictive engine into a verifiable tool for discovery.

This need for transparency is especially acute in cross-modality image-to-image translation, where the complex mapping between disparate physical domains, such as in silico labeling, often lacks a direct, human-interpretable logic^9^. Compounding this conceptual gap is a technical bottleneck: while interpretability is an emerging topic of research for classification models^50,40,51,52^, there are much fewer methods dedicated to image-to-image translation^53,54^. This leaves researchers dependent on ’off-the-shelf’ metrics that fail to capture the nuances of structural reconstructions.

In the specific context of in silico labeling, the absence of interpretability has effectively stalled the transition from proof-of-concept prototypes to reliable biological quantification. Our study addresses this gap by introducing Mask Interpreter, a generalized semantic visual interpretability method that uncovers the "explanation signatures" driving organelle predictions. We reveal distinct organelle-specific stereotypical bright-field image contexts that are required for accurate localization, demonstrating that deep learning models can capture reproducible structural biological signatures rather than latching onto spurious correlations. Furthermore, our method provides a fine-grained reliability assessment at single-cell resolution. Together with Mask Interpreter’s visual explanations, these capabilities enable the oversight required to identify when and where a model can, and cannot, be trusted, even without ground-truth fluorescence. Crucially, by leveraging these confidence scores for the automated exclusion of underperforming cells, Mask Interpreter ensures that downstream biological analyses are grounded exclusively in high-quality in silico labeling, finally allowing these models to be applied robustly to new experimental conditions.

While identifying where a model fails is a critical prerequisite for biological discovery, current methods primarily rely on quantifying the statistical stability of the generative process to flag potential errors^32,55–58^. These approaches act as diagnostic tools to flag likely errors, but they provide a statistical signal of instability rather than a structural explanation of the features that led to the failure. In contrast, Mask Interpreter provides a semantic layer of evidence by pinpointing the specific bright-field features essential for each organelle reconstruction. This allows distinguishing authentic biological signals from “stable hallucinations”, where a model confidently but incorrectly predicts a structure. From a practical standpoint, Mask Interpreter provides this visual interpretability post-hoc without requiring ensembles or multiple inference passes, reducing the computational barriers typically associated with the transition from black-box prediction to human-interpretable explanation. By identifying the ’explanation signatures’ required for accurate localization, Mask Interpreter establishes a more rigorous standard for oversight, ensuring that automated discovery is not just statistically stable but biologically grounded.

Nonetheless, the method is not without constraints. First, unlike statistical stability metrics that are computed during or after inference, Mask Interpreter requires additional training phases, introducing a computational overhead that must be balanced against the need for transparency. Second, while mask efficacy provides a robust, unsupervised global assessment of mask quality without requiring fluorescent labels, extending this reliability assessment to fine-grained single-cell scoring requires ground-truth supervision during training. While this supervised confidence modeling depends on the quality of the underlying importance masks and requires paired ground-truth data, it enables the high-resolution single-cell filtering needed for automated quality control. As a partial remedy, we demonstrate that mask efficacy can be evaluated in an unsupervised manner by measuring the degradation in prediction accuracy upon masking essential input features, providing a robust global assessment of mask quality without requiring fluorescent labels. Third, our single-cell semantic confidence validation was restricted to three organelles for which the segmentation pipeline produced reliable ground-truth masks.

Extending automated single-cell exclusion to organelles lacking a tractable segmentation-based readout remains to be demonstrated. Fourth and last, while the method provides visual clarity, the translation of these importance masks into quantitative biological rules still relies on expert oversight to contextualize semantic patterns within known cell biology.

Beyond in silico labeling, the principles underlying Mask Interpreter can be adapted to other image-to-image translation tasks in microscopy and beyond. This includes diverse applications such as automated organelle segmentation in electron microscopy^59^, image restoration and super-resolution^55,60–63^, channel multiplexing^58,64^, and cross-modality transformations between disparate imaging platforms^12,65–72^. By providing a method for enhancing trust across these deep learning-based biomedical imaging tasks, Mask Interpreter offers a broadly applicable solution for establishing transparency in complex visual reconstructions. Furthermore, these same principles can be extended to classification tasks, where an explanation signature is defined by the degradation of classification performance upon feature masking. Beyond interpretability and single-cell quality control, Mask Interpreter can serve as a diagnostic tool to guide model development: identifying the specific image features where a model fails to generate a meaningful explanation signature allows for the targeted refinement of training samples. To facilitate widespread adoption, we provide Mask Interpreter as an open-source tool that can be integrated directly into automated bioimage analysis pipelines as a real-time quality control layer. By automatically flagging or filtering predictions that lack a coherent structural explanation, our approach ensures that only high-fidelity reconstructions are passed to downstream biological quantification. Ultimately, by bridging the gap between model opacity and biologically meaningful features, Mask Interpreter transforms in silico labeling from a black-box visualization tool into a rigorous, evidence-based instrument for scientific discovery.

## Declaration of generative AI and AI-assisted technologies in the writing process

During the preparation of this work the authors used Gemini, and chat GPT in order to edit and refine the text. ChatGPT was used to design Fig. 6A. After using these services, the authors reviewed and edited the content as needed and take full responsibility for the content of the manuscript.

## Methods

### Data

We used the AICS WTC-11 hiPS cell Single-Cell Image Dataset v1^38^. From the FOV spinning-disk confocal microscopy section, we used the 16-bit z stack images, acquired with a 100× objective, with a resolution of 624 × 924 pixels and a physical pixel size of 0.108 µm × 0.108 µm. Each z stack consisted of 50–75 slices with resolution of 0.29 µm between consecutive slices. Specifically, we used the brightfield channel, DNA channel, and plasma membrane channel, for all available organelles, and the EGFP-tagged cellular structure channel for the following eight organelles: nucleoli, nuclear envelope, microtubules, actin filaments, mitochondria, endoplasmic reticulum, actomyosin bundles and Golgi. For each organelle we used a single FOV from each available well, for a total of 75-190 FOVs per organelle (average = 94.55) that were divided to 90% for train and validation and 10% for test.

We used the FOV cell segmentation label maps to extract individual cells from the FOV images. We excluded cells that were not entirely inside the FOV owing to the lack of metadata and inability to reliably extract shape descriptors. For each cell, we constructed a 3D bounding box and applied the segmentation masks to the brightfield and EGFP FOV images.

For the analysis of batch effects detection, we used metadata fields: ‘CellLineId’, and ‘Workflow’. ‘CellLineId’ corresponds to the catalog number in the Allen Cell Catalog (https://www.allencell.org/cell-catalog.html). ‘Workflow’ describes image acquisition protocol (i.e., pipeline). Pipelines 4.0, 4.1, 4.2 acquired using a broadband halogen light source and a single-camera system, while pipeline 4.4 acquired using a far-red LED light source and a 2-camera system.

### Data preprocessing

Due to the large size of the input volumes, training was conducted on image patches of size 32×128×128 voxels (z = 12.156 *μ*m, x = 13.824 *μ*m, y = 13.824 *μ*m, correspondingly) randomly cropped from the input field of views’ volumes. To ensure consistent intensity scaling voxels intensities were z-score normalized in the context of their FOV. Thus, the normalized voxel *X_i_*′ was calculated as:

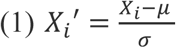

where *X_i_* is the original voxel intensity, μ is the mean voxels intensity of the FOV, σ is the standard deviation of the voxels intensities of the FOV, and *X_i_*′ is the normalized voxel intensity.

### Reproduction of Ounkomol el al.’s results

We selected ten organelles that exhibit diverse structures, cellular localizations, and in silico labeling results (Table S1) and replicated Ounkomol et al.^7^ data preprocessing, in silico labeling training of a U-Net architecture, and evaluation. The in silico labeling models we trained achieved results similar to those reported in Ounkomol et al.^7^ (Fig. S1), with minor differences attributable to random variations in data splitting between training and test sets, different patch sizes (32x64x64 pixels in Ounkomol et al.^7^ versus 32x128x128 pixels here), and U-Net implementation (Pytorch in Ounkomol et al.^7^ versus Tensorflow here).

### In silico labeling of a field of view

Similarly to Ounkomol et al.^7^, in silico labeling was performed on overlapping patches that were merged using a weighted triangle function *w*(*u*), assigning more weight to the predictions at the center of each patch and less weight to those at the peripheral areas:

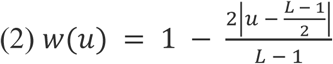

where *u* is the coordinate within the patch along a given x-, y-, or z-axis, and *L* is the size of the patch along that axis. The weights peak at the center of the patch and decrease linearly towards its boundaries. The overall weight *W*(*x*, *y*, *z*) at voxel (*x*, *y*, *z*) is calculated as the product of the weights along each dimension:

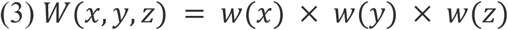

The final in silico labeling at each voxel was set by averaging the weighted predictions from all overlapping patches.

### Mask Interpreter’s implementation

#### Network architecture

Mask Interpreter optimizes an image-to-image translation model that maps a bright-field image and its corresponding in silico prediction to an importance mask, using a pre-trained in silico labeling model that remains frozen throughout training (Fig. 1C-F), the use of the in silico prediction as an input serves as an initializer for the optimization of the importance mask. Mask Interpreter employs a three-dimensional U-Net architecture consisting of four convolutional blocks in the encoder (downsampling path) with 32, 64, 128, and 256 filters, respectively, and four convolutional blocks in the decoder (upsampling path) with 256, 128, 64, and 32 filters, respectively. Each convolutional block includes a 3D convolutional layer, batch normalization, and a rectified linear unit (ReLU) activation function. Transposed 3D convolutional layers were used in the decoder blocks for upsampling. Mask interpreter optimizes an importance mask, in the size of input image patch, with voxel values ranging between 0 and 1 using a sigmoid activation function in the final layer.

### Setting the noise parameter

Gaussian noise was added to the original bright-field image in a manner modulated by the importance mask, during Mask Interpreter’ training. Specifically, the noisy image *I_i_*′ at each voxels *i* was computed using the equation:

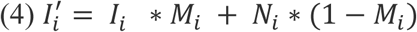

where *I_i_* is the original normalized bright-field voxel intensity, *M_i_* is the corresponding importance mask value, and *N_i_* is a noise value sampled from a Gaussian distribution with mean zero and a standard deviation σ. To determine σ we sampled 10 bright-field FOVs for each organelle’s in silico labeling model. Gaussian noise with varying σ values ranging from 0.0 to 4.0 standard deviations were added to each voxel in these images, and the Pearson correlation coefficient (PCC) between the in silico labeling predictions with and without the added noise was measured. For each in silico labeling model we selected the lowest σ for which the PCC dropped below 0.2, indicating a major degradation in prediction performance due to the introduction of noise (Fig. S15). Choosing a much lower σ would fail to reveal regions where noise can hamper the in silico labeling prediction.

### Loss terms and optimization

The Mask Interpreter model was trained using backpropagation to optimize a composite loss function consisting of three terms: target score loss, mask minimization loss, and similarity maximization loss. These loss terms work in tandem to ensure that the importance mask highlights essential voxels in the bright-field image for the fixed in silico labeling model, in the sense that inclusion of noise in these voxels will lead to deteriorated in silico labeling. The primary objective was to maximize the inclusion of noise to non-essential voxels while ensuring that the fixed in silico labeling prediction from the noisy bright-field image *Ŷ*′ remains highly similar to the prediction from the original unperturbed bright-field image *Ŷ*.

The target score loss term enforced this similarity by maintaining a PCC of at least 0.9 between *Ŷ*′, *Ŷ*. More formally:

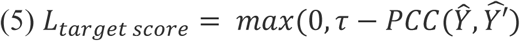

Where the parameter τ defines a threshold that controls the trade-off between the amount of noise incorporated to the bright-field image and the in silico labeling fidelity. The target score loss is positive when the PCC drops below τ, penalizing the model upon deteriorated in silico labeling from the noisy bright-field image. Namely, τ→1.0 severely limits the inclusion of noise - inducing uninformative “everything matters” masks, whereas a low τ allows excessive noise inclusion breaking the in silico labeling fidelity. This definition of the target score loss enables users to set τ in a human interpretable manner. Empirically, we found that τ = 0.9 balanced the generation of informative masks with acceptably similar predictions structure.

The mask minimization loss term maximized the inclusion of noise to the bright-field image by minimizing the sum of the importance mask values, enforcing sparseness:

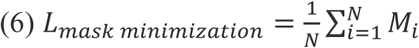

Where *M_i_* is the value of the importance mask at voxel i and *N* is the total number of voxels in the mask. Minimizing the mask minimization loss enforces the reduction of voxels deemed as “important”, thus allowing the inclusion of more noise to voxels that can be perturbed without causing major alterations for the in silico labeling.

The similarity maximization loss term ensures that the in silico labeling voxels’ intensity remain similar following the inclusion of noise in terms of the mean squared error (MSE):

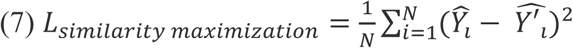

Where *Ŷ_i_*′, *Ŷ_i_* are the in silico predictions at voxel i from the unperturbed and from the noisy bright-field images, respectively. *L_target score_* and *L_similarity maximization_* are complementary, where the former enforces structural fidelity and the latter enforces intensity fidelity.

The overall loss for training a Mask Interpreter model was the weighted sum of the three loss terms:

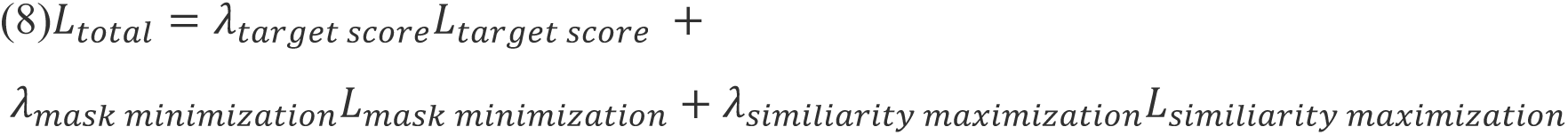

We empirically set λ_target score_ = 6, λ_mask minimization_= 1, and λ_similarity maximization_=1, defining the target score loss as the most dominant term. Once the target score constraint was satisfied, *L_target score_* effectively reduced to zero and the optimization shifted toward balancing *L_mask minimization_* and *L_similiarity maximization_*. A systematic ablation analysis confirmed that all loss terms were necessary (Fig. S2). Mask Interpreter training was executed on Nvidia RTX3090 GPUs, with organelle-specific training times ranging from 6 to 12 hours on a single GPU.

### Generation of explanation signatures

The trained Mask Interpreter U-Net receives the original bright-field image and its in silico prediction and generates the corresponding importance mask. An organelle-specific threshold was set to binarize the importance mask into an explanation signature as follows. Candidate thresholds were evenly sampled between 0.0 and 1.0.

For each candidate threshold, and for each FOV in test set, the importance mask was binarized and used to introduce noise to the bright-field image: voxels falling below the threshold were considered non-essential and subjected to noise, while voxels above the threshold were preserved as is. The Pearson correlation coefficient (PCC) between the predictions derived from the noisy and unperturbed bright-field images was computed to quantify the “mask efficacy”. An organelle-specific threshold was empirically set as the smallest value for which the mean mask efficacy fell below 0.87. This procedure ensures that the binary explanation signature balances minimal coverage of voxels deemed “important”, i.e., sparser explanation signatures, with the maintenance of high predictive fidelity from the derived noisy bright-field image. Fig. S16 depicts mask efficacy as a function of the explanation signature threshold for the ten organelles presented in this study.

### Visualization of a representative focal plane

For each field of view presented in this manuscript, we manually selected a single z-slice that best captured the cells in focus to ensure optimal visual clarity.

### Implementation and benchmarking of gradient-based alternative explanation methods for image-to-image translation

We adapted GradCAM^40^ and Guided Backpropagation^41^, originally developed for image classification, to our in silico labeling setting by following a procedure similar to the “SEG-GRAD-CAM” approach described in Vinogradova et al^53^. First, instead of focusing on a single class logit as in classification models, we treated each organelle’s fluorescence prediction voxel as an equivalent “class score” in the U-Net. For GradCAM, we targeted the final encoder layer (the bottleneck of the 3D U-Net) and computed the gradients of the predicted organelle signals with respect to the corresponding feature maps. Following Vinogradova et al., we aggregated these gradients to determine channel-wise importance weights and generated heatmaps by taking the weighted sum of the activation maps. A ReLU function was then applied to emphasize regions that contribute positively to the prediction. For Guided Backpropagation, we computed partial derivatives of the predicted fluorescence signal with respect to the bright-field input intensities, applying the same modifications described by Springenberg et al^50^.

To systematically benchmark Mask Interpreter versus GradCAM and Guided Backpropagation we conducted a noise perturbation test. For each noise level, from 0% to 100%, and for each FOV, we used binary search to identify a threshold that would yield the specified fraction of voxels in the corresponding importance mask/gradient map. We then applied Gaussian noise exclusively to those voxels in the bright-field image and performed in silico labeling inference. This procedure was repeated for each FOV from the test set (up to 10 FOVs) for each organelle-specific model. The resulting mask efficacy between the noisy and unperturbed predictions were plotted against the percentage of noised voxels (Fig. 3E), allowing a direct comparison of how well each explainability method preserves prediction accuracy under increasing noise levels.

### Quantification of in silico labeling results of Mitotic vs Interphase cell state

We analyzed 16 and 11 test set FOVs for the nuclear envelope and microtubules, respectively. These two organelles are known to change their structure and organization during mitosis in a manner hampering their in silico labeling^31^ and was used here to demonstrate association between non-sterotypical explanation signatures and deteriorated in silico labeling. For each single cell we used its accompanied segmentation to generate a cell-specific binary mask that isolated the cell from its neighbors. This mask was then element-wise multiplied with both the fluorescence (ground truth) and in silico prediction channels. The resulting volumes were cropped to the 3D bounding box of the mask to create tightly bounded single-cell images. Cells intersecting FOV boundaries were excluded due to missing metadata in the WTC-11 dataset. Using the cell_stage metadata cells were categorized into interphase or mitotic stages. This procedure yielded approximately 160 single cells per organelle. The Pearson correlation coefficient (PCC) was subsequently calculated between the masked prediction and ground truth voxels.

### Spatial PCC visualization

Spatial PCC visualization (e.g., Fig. 4D) was computed by partitioning the FOV using a sliding window of size 64×64x16 pixels in the x, y, and z dimensions, respectively, with an overlap of 16 × 16 × 1 pixels.

Spatial PCC was computed between the in silico labeled and ground truth voxel intensities for each window and averaged for each voxel across all the overlapping windows that involved that voxel.

### Detection of batch effects

We compiled 1,821 FOVs pooled across 18 imaging batches, where an ’imaging batch’ was defined as each one of 18 genetically edited cell lines, where each was fluorescently tagged for a specific organelle marker and imaged under the same imaging pipeline. Mask efficacy was computed for each FOV as the PCC between the in silico predictions obtained from the unperturbed and from the noisy bright-field image. We identified an imaging pipeline as a putative batch effect when the corresponding FOVs consistency reported low mask efficacies compared to held-out test FOVs of the training batch (e.g., Fig. 5D). A putative batch effect was verified according to its FOVs’ in silico labeling performance (e.g., Fig. 5E). Statistical significance was assessed at the FOV level, for each organelle model and imaging pipeline. Values from each pipeline were compared with the corresponding organelle-specific imaging batch test set distribution using a one-sided Mann-Whitney U test. The null hypothesis was that the pipeline distribution was not shifted toward lower values than the test set distribution. Rejection of this null hypothesis indicated that FOVs acquired with the tested pipeline had significantly lower mask efficacy or in silico labeling performance than the test set control.

### Density Resampling Estimate of Mutual Information (DREMI)

The joint distribution of the mask efficacy and the in silico labeling concentrate in a narrow range of values (e.g., Fig. 5C) and thus obscure the true dependency between these parameters across their full dynamic range. To remedy this limitation we applied Density Resampled estimate of Mutual Information, DREMI^44^, which estimates the in silico labeling distribution conditioned on the partitioning of mask efficacies to evenly spaced bins. DREMI was computed using kernel density estimation (KDE) method to derive joint and conditional probability distributions^44^

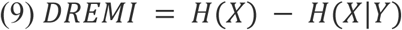

where *H*(*X*) represents the marginal entropy, and *H*(*X*|*Y*) is the conditional entropy of the in silico labeling accuracy given the mask efficacy score.

### Supervised quality assessment

#### Architecture and training of the confidence models

We finetuned the 3D ResNet-18 architecture ‘r3d_18’^73^, that was pretrained on the Kinetics Human Action Video Dataset^74^. This architecture was selected because it was suitable for 3D images, showed good results on medical image segmentation tasks^75^, and because similar approaches succeeded in related tasks such as assessing segmentation quality control^76^. The model was adapted for two-channel input of the in silico labeling and its corresponding importance mask, by adapting the initial convolutional layer to accept 2-channel volumetric data. The final classification head was replaced with a regression module comprising two fully connected layers (512→128→1), with ReLU activations. The model outputs a scalar between 0 and 1 that is the estimated error of the in silico labeling prediction in terms of the absolute PCC deviation between the in silico labeling prediction and fluorescent ground truth. During training, the confidence model was optimized to minimize the Mean Squared Error (MSE) between the true and the predicted error, defined as 1 - PCC between the predicted patch and the original fluorescence image. Three fine-tuning strategies were evaluated: (1) training only the added regression layers. (2) fine-tuning the final residual stage of the model’s backbone together with the regression layers. (3) training the entire model. In the partial fine-tuning configuration (configuration 2), all parameters in the fourth residual stage (layer4) and the added fully connected layers (fc1 and fc2) were trained, whereas the stem and the first three residual stages (layer1–layer3) were frozen. Partial fine-tuning yielded the best tradeoff between performance and generalization. Fig. 6A provides a schematic of the confidence model pipeline.

### Data partitioning for confidence model training

Since the confidence model is trained using outputs generated by the in silico labeling model and by the Mask Interpreter model, it is essential to maintain consistent train-test splits such that FOVs that were used for evaluating these models are not exposed to the confidence model during training. Thus, we preserved the original Mask Interpreter models’ train-test partitioning to prevent data leakage. Train-test partitioning was performed at the FOV level, and patches were subsequently extracted from each FOV to train the confidence model as explained above.

### Confidence model evaluation

We evaluated the supervised confidence model using metrics that measure the difference between the predicted and the true in silico labeling errors, namely Mean Squared Error, Pearson Correlation Coefficient and R^2^. In addition, we performed an ablation experiment to confirm that the inclusion of the importance mask contributes to the model by comparing it to a single-channel confidence model that predicts the in silico labeling error using the in silico labeling prediction. To directly compare two confidence measurements or models, we used a paired model-comparison analysis. For each image patch *i* and confidence measure *m*, we calculated the absolute error between the estimated in silico labeling performance and the measured performance:

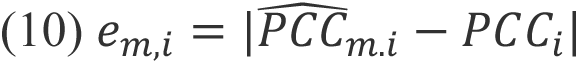

where 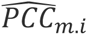 is the PCC estimated by confidence measure *m*, and *PCC_i_* is the measured Pearson correlation between the in silico labeling prediction and the corresponding fluorescence image. We then compared two confidence measures by calculating:

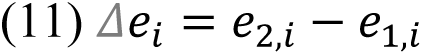

### Threshold selection for classifying low confidence in silico labeling regions

We manually selected a per-FOV threshold to visualize low-quality predicted and ground-truth-verified in silico labeling regions (e.g., Fig. 6C). The thresholds were selected to highlight low-confidence regions while avoiding excessive background or scattered noise.

### Extracting single-cells from FOVs

We adapted the analysis to predict the in silico labeling performance at single cell resolution. First, we followed our FOV-scale standard in silico labeling inference pipeline: generating volumetric predictions from overlapping 3D patches, and then merging them into a full FOV prediction using weighted averaging with triangular weights (as described above). Next, we extracted individual cell masks within each FOV (as described above) and used those masks on the in silico labeling, fluorescence ground truth, explanation masks generated by the Mask Interpreter and confidence scores.

### Organelle segmentation quality control analysis

Organelle segmentation followed the pipelines provided by the Allen Institute for Cell Science through their Structure Segmenter^77^. The fluorescence ground-truth segmentations were available as part of the Allen Institute’s dataset. Organelles chosen for this analysis were the nuclear envelope, the nucleoli and the microtubules, for which the Allen Institute segmentation pipeline produced reliable ground-truth masks. To quantify model performance at the single-cell level, we evaluated predicted organelle labels within segmented regions of interest (ROIs). Each ROI corresponded to an individual cell, and for every cell we computed overlap-based segmentation metrics between the predicted segmentation and the corresponding ground-truth segmentation. Because the 3D fluorescence signal of a cell is not uniformly informative across the z-axis, we first identified the 16 most informative slices per cell. These “peak-signal slices” were defined as the contiguous z-window with the highest integrated fluorescence intensity within the cell mask. This approach focused the evaluation on the region of the volume in which the organelle structure was most visible and well resolved, while avoiding planes with little or no relevant signal, such as out-of-focus or empty slices. For each metric, performance was computed across these 16 slices and summarized as the mean value per cell.

Using these cell-level metrics, including IoU and Dice, we examined whether predicted confidence stratified downstream segmentation quality. For each organelle, cells were divided into confidence quartiles, with Q1 corresponding to the lowest-confidence cells and Q4 to the highest-confidence cells. To test whether low confidence identified poor segmentation performance, metric values in Q1 were compared with those in the pooled Q2–Q4 group using a one-sided Mann–Whitney U test^78^, with the alternative hypothesis that segmentation performance was higher in Q2–Q4. Cliff’s δ^79^ was calculated as a non-parametric measure of effect size. Because one prespecified comparison was performed within each organelle, no within-organelle multiple-comparison correction was applied.

## Data and Code availability

Mask Interpreter source code is publicly available on GitHub. The core implementation is available at https://github.com/zaritskylab/MaskInterpreter. A companion repository, https://github.com/zaritskylab/MaskInterpreter-Applications, provides the supervised confidence model implementation and the single-cell Mask Interpreter application workflow. The trained Tensorflow in silico labeling and Mask Interpreter models, example data, and the train-test split files are available at Zenodo: https://zenodo.org/records/18590674. The trained supervised confidence models and precomputed result arrays, together with the models and example data required for the single-cell Mask Interpreter workflow, are available at https://doi.org/10.5281/zenodo.20522083.

## Funding and Acknowledgments

This research was supported by the Israel Science Foundation (ISF, grant No. 2516/21, to A.Z), by the Israel Council for Higher Education (CHE) via the Data Science Research Center, Ben-Gurion University of the Negev, Israel (to A.Z), and by the Allen Family Philanthropies (to A.Z). This research was supported by the Allen Institute, founded by Jody Allen – chair and co-founder of Allen Family Philanthropies, and the late Paul G. Allen – investor, philanthropist, and co-founder of Microsoft. We gratefully acknowledge their vision and generosity, which make this work possible. We thank Noga Levy for critically reading the manuscript. We thank Orit Kliper-Gross for critically reviewing our GitHub repository.

## Author Contribution Statement

L.B.N. and A.Z. conceived the study. L.B.N. developed the computational method and analyzed the data for Mask Interpreter. G.M. developed the computational method and analyzed the data for the confidence model.

N.E. performed analysis. L.B.N., G.M. and A.Z. interpreted the data and drafted the manuscript. M.P.V., J.C., N.G., and S.M.R. provided feedback, reviewed and revised the manuscript. A.Z. mentored L.B.N., G.M., and N.E. All authors edited the manuscript and approved its content.

## Competing Interests Statement

L.B.N. G.M. and A.Z. are inventors on a US Provisional Application, number 64/124,279 filed by B. G. Negev Technologies and Applications, which covers the methods described in this manuscript.

## Supplementary Information

### Supplementary Figures

**Figure S1.**
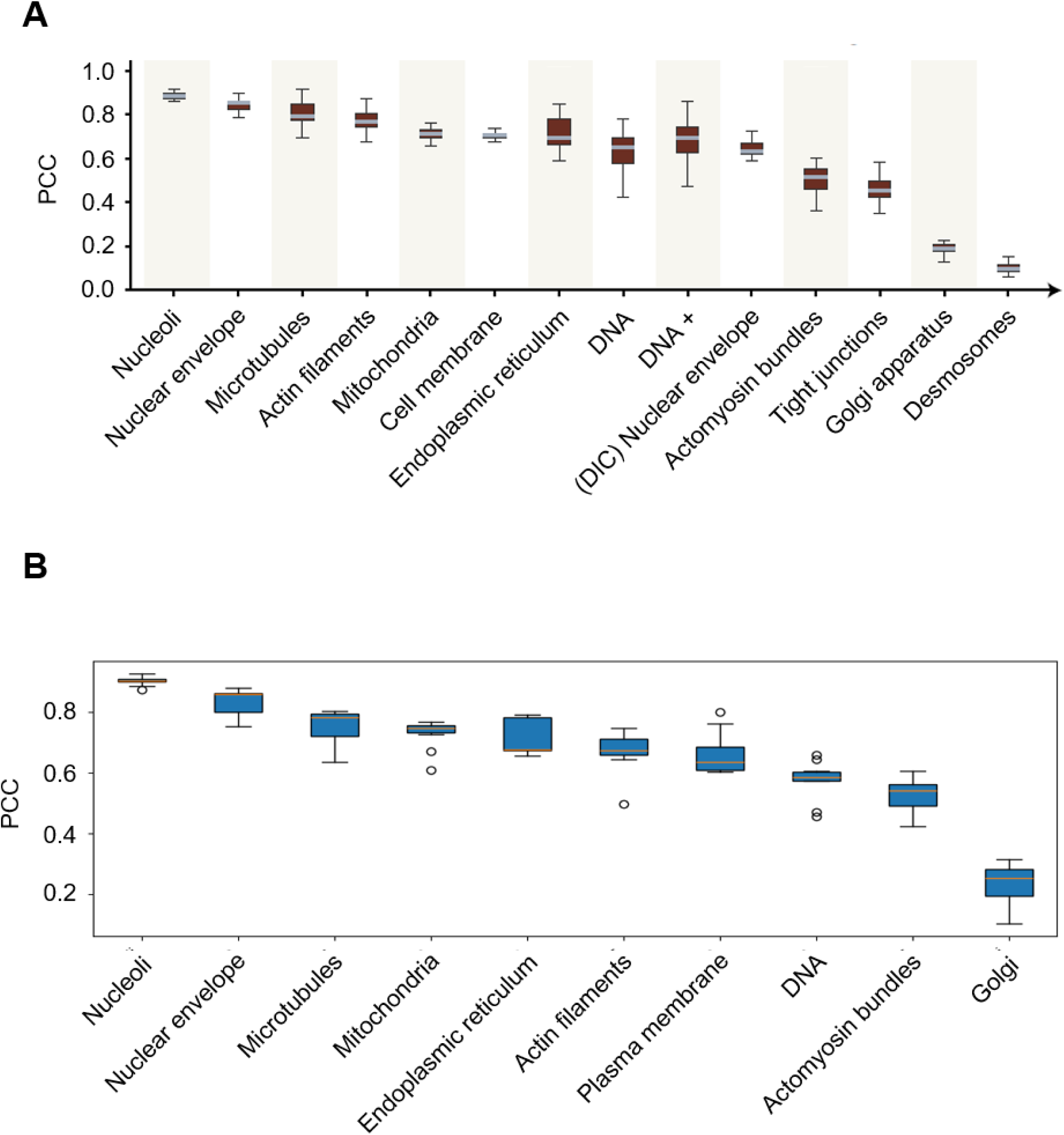
Reproduction of the in silico labeling results reported in Ounkomol et al.(A-B) Box plots show the median and interquartile range (IQR); whiskers extend to the most extreme values within 1.5 × IQR. Outliers are shown as white circles only on bottom panel **(A)** Distribution of the field-of-view-level Pearson correlation coefficients (PCC) between fluorescence ground truth and in silico labeling predictions across subcellular structures), generated using data from Ounkomol et al.^7^. Adapted with permission from Springer Nature: Nature Methods (Ounkomol et al., 2018). **(B)** In silico labeling replication results across 10 organelles (Methods). The range of test FOVs for each organelle was 8-14. Note, that the specific images used in A were not reported in Ounkomol et al.^7^ hampering the possibility for using the exact same images for replication.

**Figure S2.**
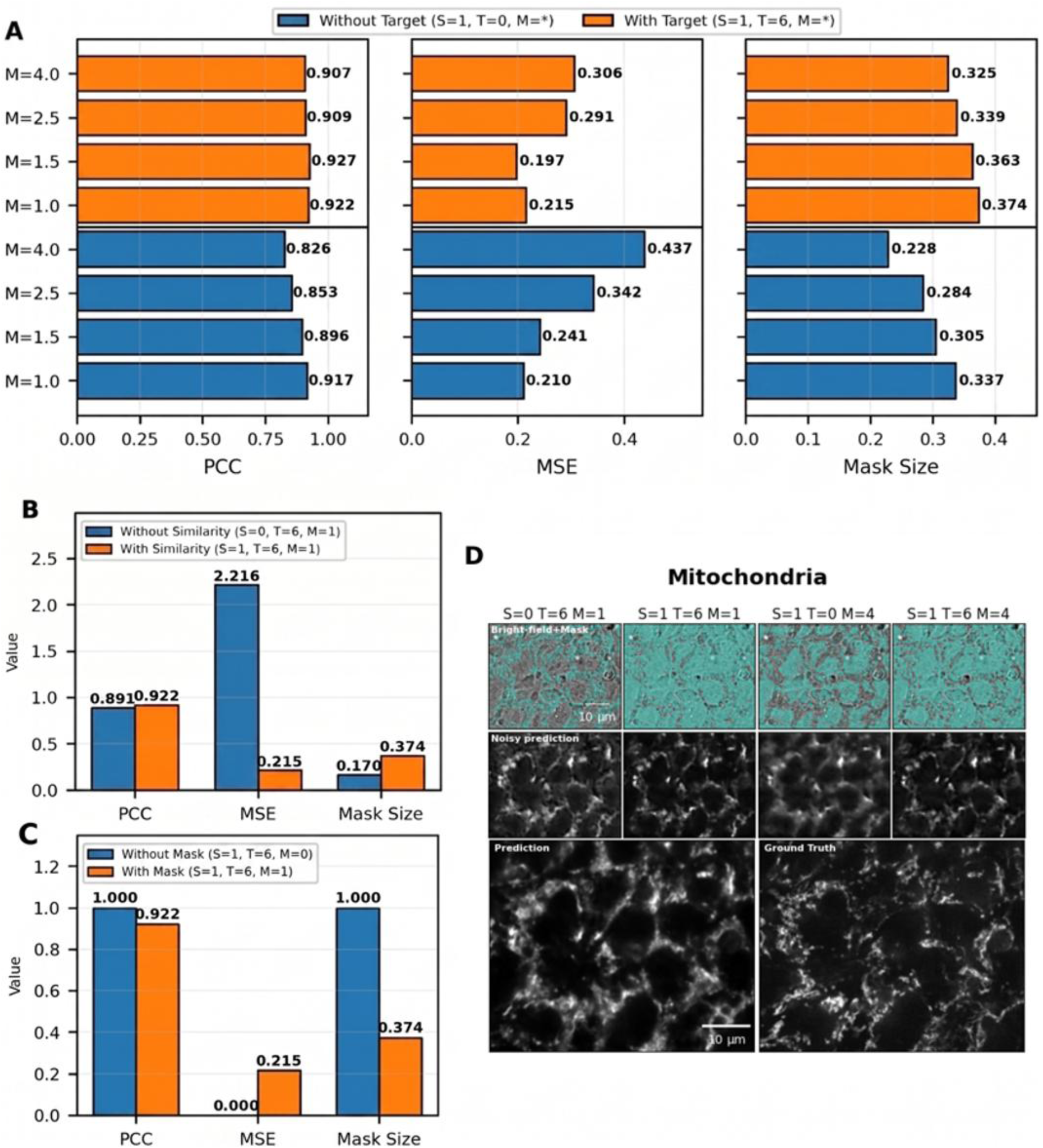
Ablation analysis. (**A–C**) Quantitative evaluation of the contribution of each of the three loss terms to a Mask Interpreter model trained to explain the mitochondria in silico labeling model. The weights for each loss term are marked with a letter: target score with ‘T’, similarity maximization with ‘S’, and mask minimization with ‘M’. Each ablation experiment trained a Mask Interpreter model after deviating one of the loss terms’ weights, while fixing the weights of the other loss terms. We selected the mitochondria for the ablation study because its in silico model used vast spatial context around the organelle (see Fig. 2B) and thus we hypothesized that different S,T, and M weights will alter the mask interpreter outcome. Three metrics were evaluated for each loss weights configuration: (1) Structural fidelity. The Pearson correlation coefficient (PCC) between the in silico predictions from the noisy and the unperturbed bright-field images (higher is better), (2) Intensity consistency. The mean squared error (MSE) between the in silico predictions from the noisy and the unperturbed bright-field images (lower is better), and (3) Mask minimization. The fraction of pixels above a threshold of 0.2 in the importance mask (lower is better). Importantly, low quality in silico labeling from the noisy image (i.e., low structural fidelity and/or high intensity consistency) implies that the importance mask does not encode the bright-field voxels’ importance and thus cannot provide a reliable visual explanation. (**A**) Ablation (T = 0, blue) versus inclusion (T = 6, orange) of the target score loss term across different weights for the mask minimization loss term (M = 1, 1.5, 2.5, 4). Ablation of the target score loss reduced structural fidelity and intensity consistency but improved the mask minimization. (**B**) Ablation (S = 0, orange) versus inclusion (S = 1, blue) of the similarity maximization loss term. Ablation of the similarity maximization loss maintained structural fidelity, but degraded the intensity consistency by an order of magnitude, and improved the mask minimization. (**C**) Ablation (M = 0, blue) versus inclusion (M = 1, orange) of the mask minimization loss term. Ablation of the mask minimization loss trivially preserved perfect structural fidelity and intensity consistency, at the cost of a maximal, non-informative mask that covers the entire image. (**D**) A representative in-focus z-slice for mitochondria for different loss configurations (columns). Top row: The explanation signature (cyan) is overlaid on the bright-field image. Middle row: The corresponding in silico prediction from the noisy bright-field image (“noisy prediction”). Bottom row: In silico labeling from the unperturbed bright-field image (“prediction”), and the corresponding fluorescence ground truth. Loss configurations that violate either structural fidelity (S=1, T=0, M=4) or intensity consistency (S=0, T=6, M=1) result in degraded in silico labeling from the noisy bright-field, whereas the full loss formulation (S=1, T=6, M=1 and S=1, T=6, M=4) yields interpretable explanation signatures while preserving prediction fidelity. Scale bars = 10 μm.

**Figure S3.**
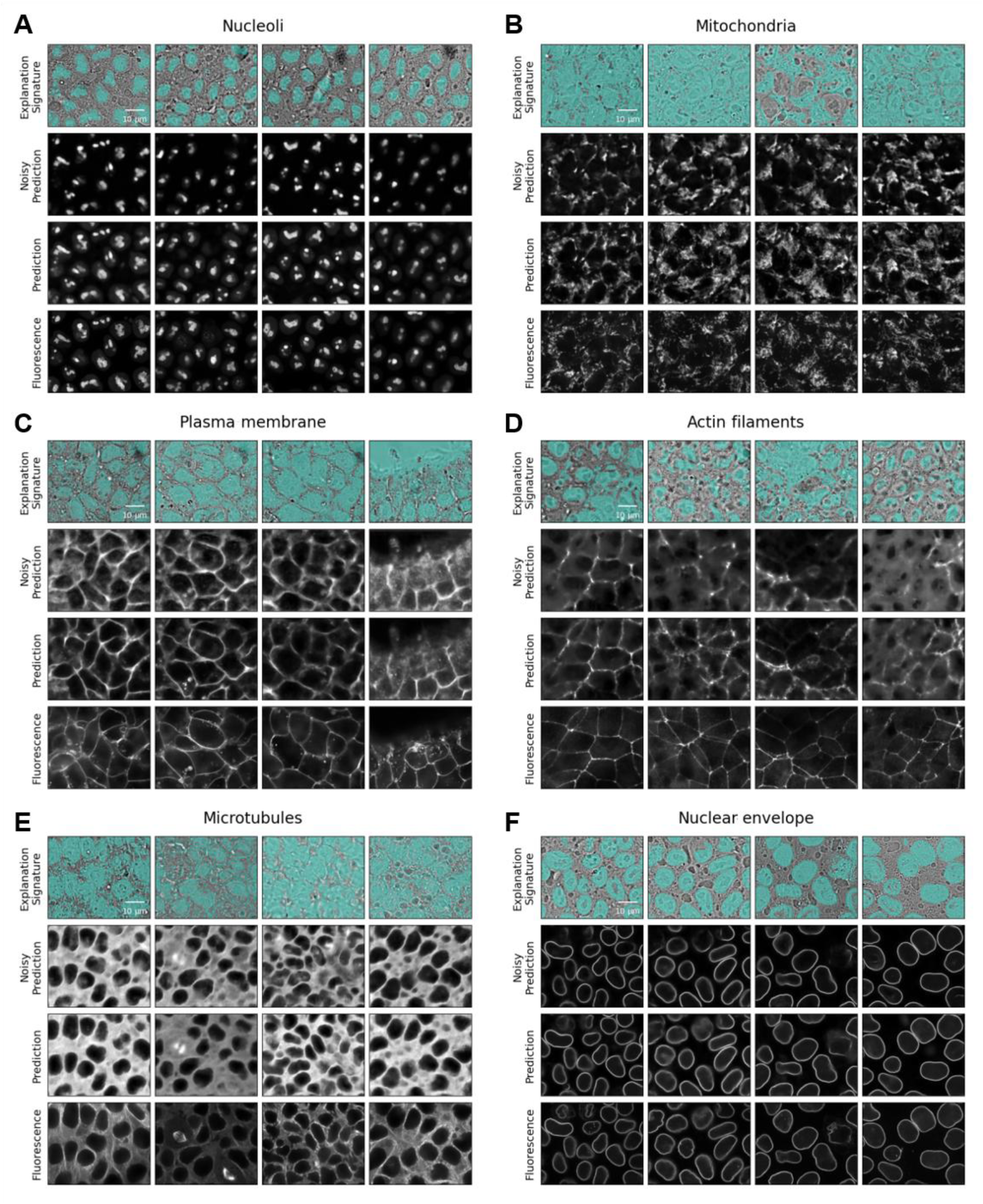
Examples of Mask Interpreter explanation signatures across organelles. Manually selected in-focus z-slices in representative fields of view (columns) demonstrating the consistency of the Mask Interpreter’s explanation signatures across six organelles. Rows display (top-to-bottom): the bright-field image overlaid with the binarized importance mask; the in silico labeling prediction from the noisy input; the prediction from the unperturbed input; the ground-truth fluorescence. Thresholds used for mask binarization: (**A**) nucleoli = 0.6, (**B**) mitochondria = 0.2, (**C**) plasma membrane = 0.4, (**D**) actin filaments = 0.2, (**E**) microtubules = 0.1, (**F**) nuclear envelope = 0.2. Scale bars = 10 μm.

**Figure S4.**
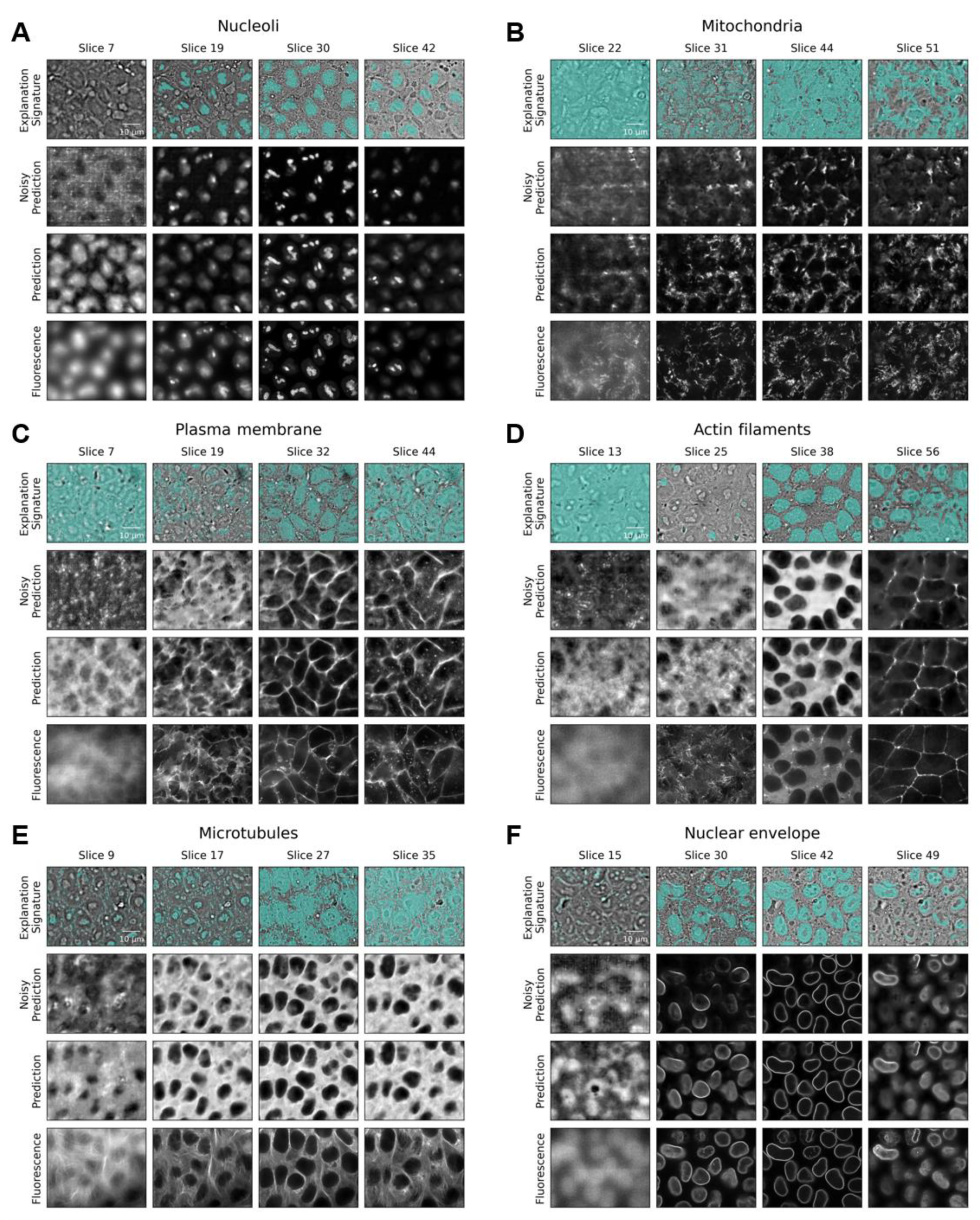
Examples of Mask Interpreter explanation signatures across organelles in 3D. Axial slices from representative 3D fields of view (columns). Rows display (top-to-bottom): the bright-field image overlaid with the binarized importance mask; the in silico labeling prediction generated from the noisy input; the prediction from the unperturbed input; the ground-truth fluorescence. Thresholds used for mask binarization: (**A**) nucleoli = 0.6, (**B**) mitochondria = 0.2, (**C**) plasma membrane = 0.4, (**D**) actin filaments = 0.2, (**E**) microtubules = 0.1, (**F**) nuclear envelope = 0.2. Scale bars = 10 μm.

**Figure S5.**
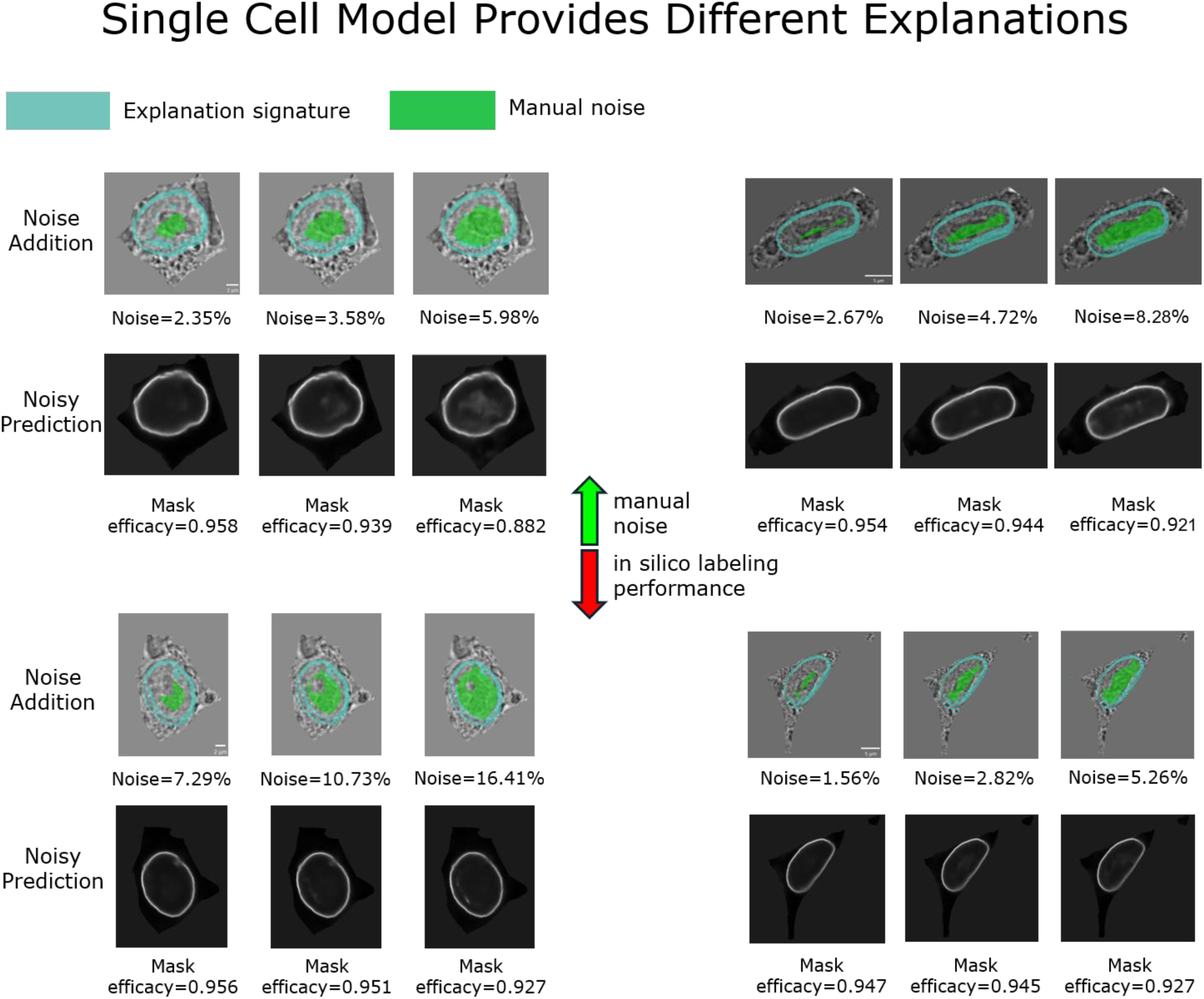
Single cell-centric in silico labeling of the nuclear envelope does not rely on the nucleus. Manual introduction of noise in the bright-field image regions corresponding to the nuclear interior of 4 representative cells did not hamper the single cell-centric in silico labeling of the nuclear envelope. Top panels: noise was manually added to the bright-field image regions corresponding to the nuclear interior (green) that did not intersect with the single cell-centric Mask Interpreter’s explanation signatures (cyan). Bottom panels: the single cell-centric in silico labeling prediction of the corresponding noisy bright-field images had a minimal effect, qualitatively and quantitatively according to the corresponding mask efficacies, defined as the Pearson correlation between the unperturbed and the noisy predictions.

**Figure S6.**
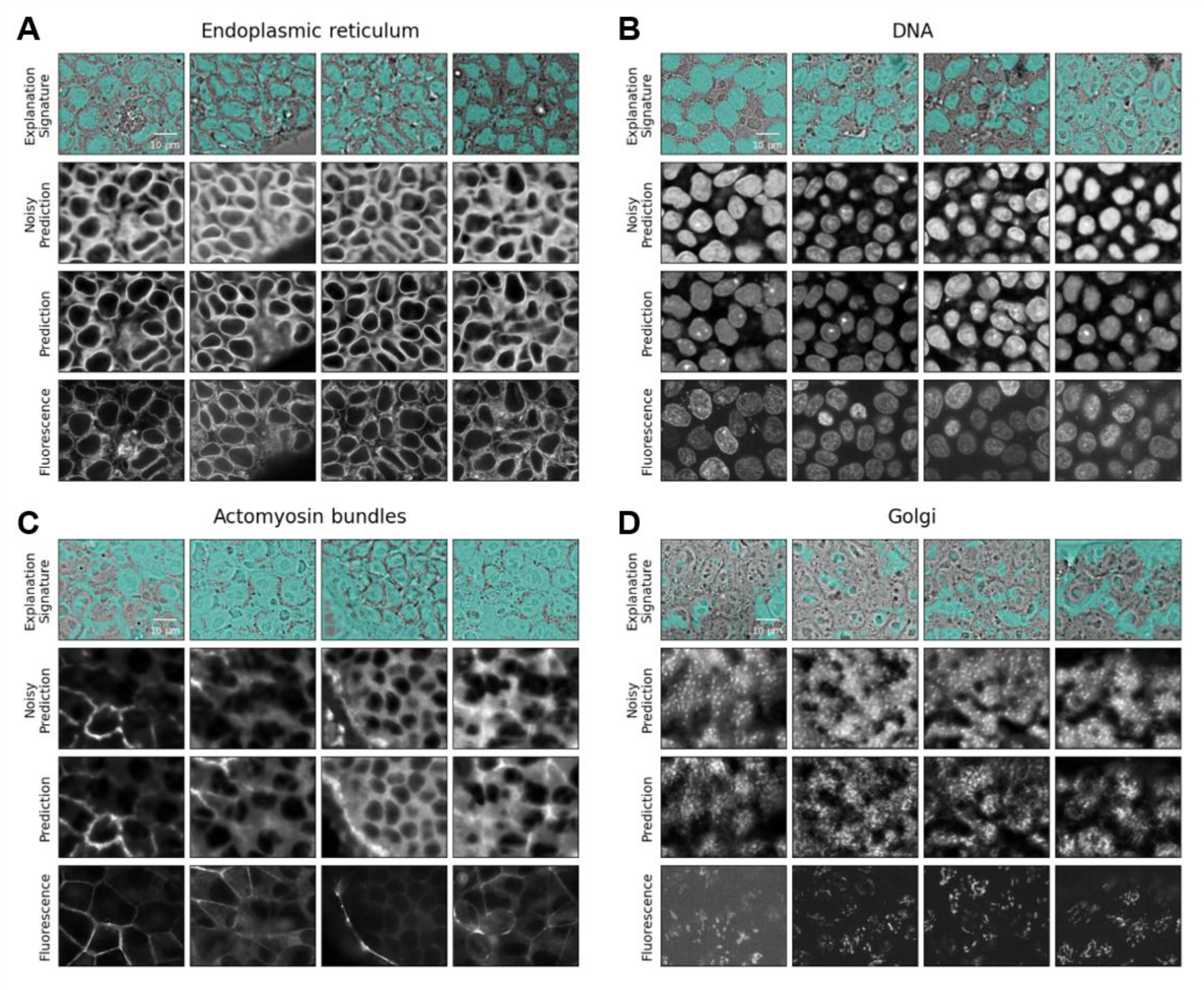
Examples of Mask Interpreter explanation signatures across organelles. Manually selected in-focus z-slices in representative fields of view (columns) demonstrating the consistency of the Mask Interpreter’s explanation signatures across four organelles. Rows display (top-to-bottom): the bright-field image overlaid with the binarized importance mask; the in silico labeling prediction from the noisy input; the prediction from the unperturbed input; the ground-truth fluorescence. Thresholds used for mask binarization: (**A**) endoplasmic reticulum = 0.4, (**B**) DNA = 0.3, (**C**) actomyosin bundles = 0.02, (**D**) Golgi = 0.3. Scale bars = 10 μm.

**Figure S7.**
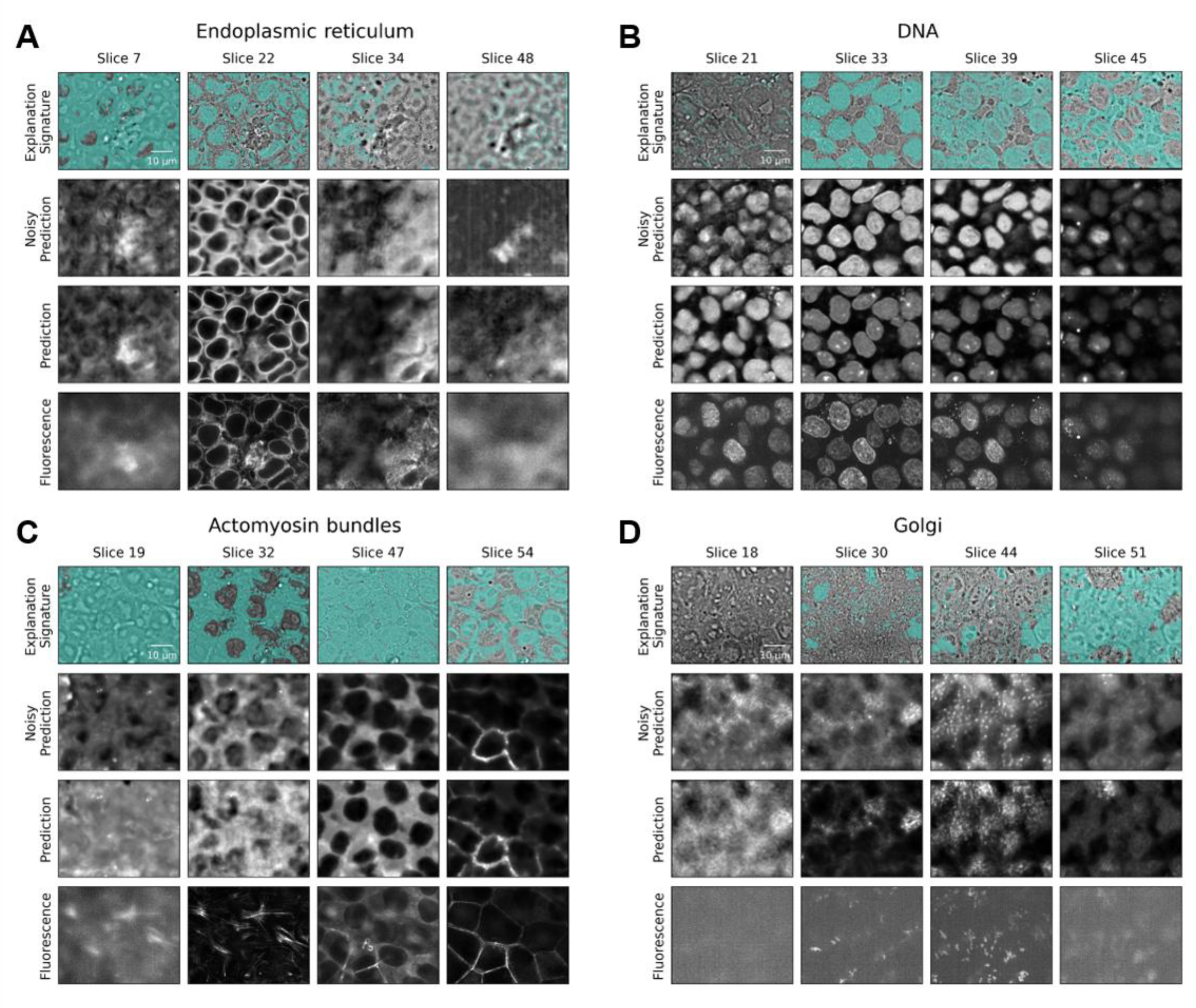
Examples of Mask Interpreter explanation signatures across organelles in 3D. Axial slices from representative 3D fields of view (columns). Rows display (top-to-bottom): the bright-field image overlaid with the binarized importance mask; the in silico labeling prediction generated from the noisy input; the prediction from the unperturbed input; the ground-truth fluorescence. Thresholds used for mask binarization: (**A**) endoplasmic reticulum = 0.4, (**B**) DNA = 0.3, (**C**) actomyosin bundles = 0.02, (**D**) Golgi = 0.3. Scale bars = 10 μm.

**Figure S8.**
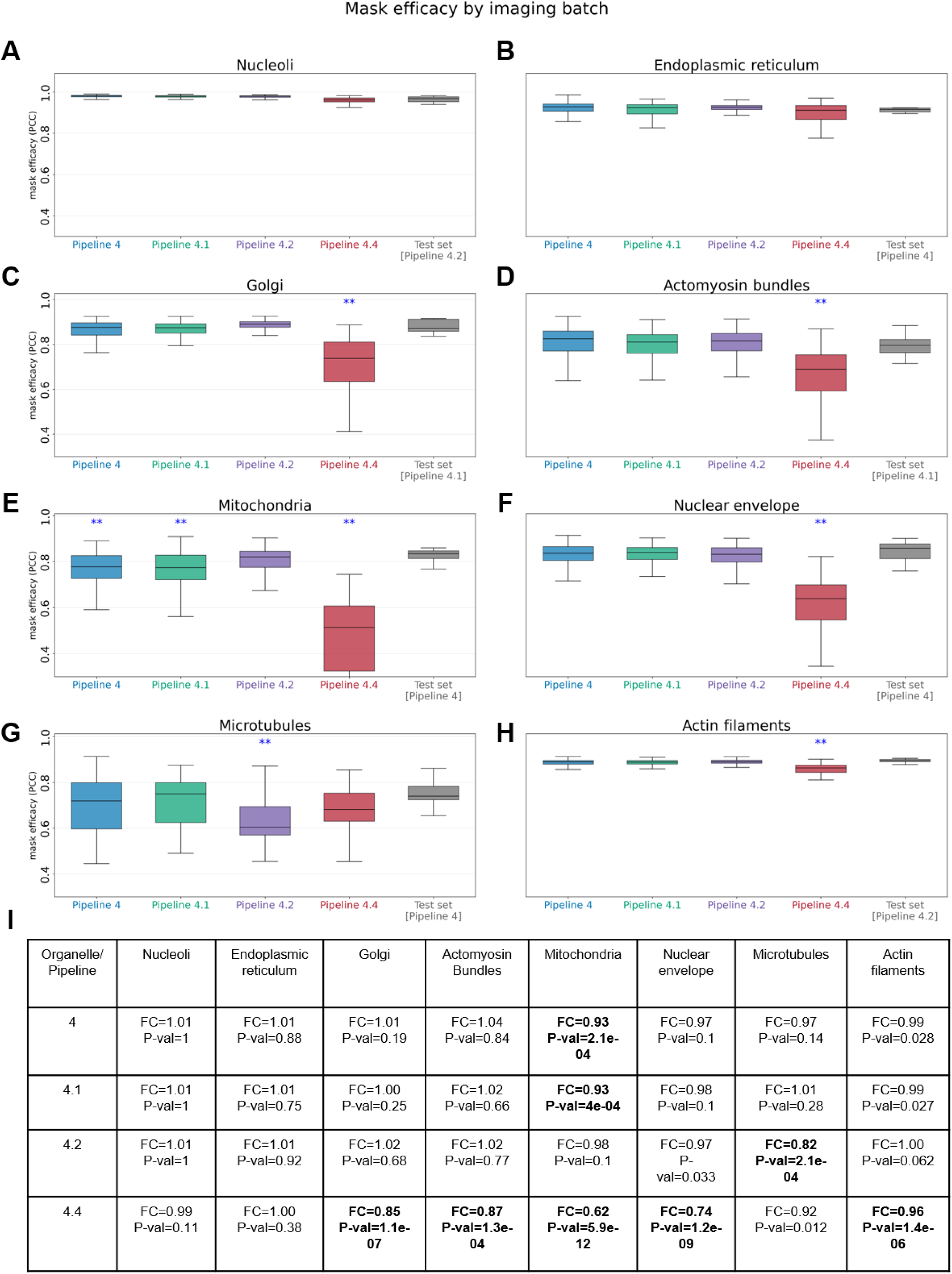
Mask Interpreter predicts batch effects in some organelles, but not others. **(A-H)** Box plots of FOVs’ mask efficacies across four imaging pipelines for eight organelles. Notably, each organelle was acquired using only one of the imaging batches by using one of four imaging pipelines and thus the fluorescence ground truth is absent. Boxes indicate the interquartile range with the median shown as the center line; whiskers extend to 1.5× the interquartile range (outliers not shown). While most pipelines exhibit comparable mask efficacy values, there’s a drop in mask efficacy for 5 out of 8 organelles for imaging pipeline 4.4, indicating a workflow-specific deviation in the bright-field input distribution. For each imaging pipeline, mask efficacy values were compared with the corresponding test set distribution of the relevant organelle using a one-sided Mann–Whitney U test, testing whether the pipeline showed lower mask efficacy than the test set. The test set was the same test set used for the Mask Interpreter and in silico labeling models. Imaging pipeline used to acquire each organelle test set FOVs is mentioned in parentheses in the x-axis. Test set controls included N = 15, 7, 8, 8, 18, 16, 11, 10 FOVs, for nucleoli, endoplasmic reticulum, Golgi, actomyosin bundles, mitochondria, nuclear envelope, microtubules, actin filaments respectively, and the different pipelines included N = 641, N = 465, N = 446, and N = 116 FOVs for pipelines 4.0, 4.1, 4.2, and 4.4, respectively, **P < 0.01. (**I**) Summary table reporting the fold changes (FC) and the corresponding *P*-values for each organelle–pipeline comparison. Statistical significant decrease in mask efficacy is marked in bold.

**Figure S9.**
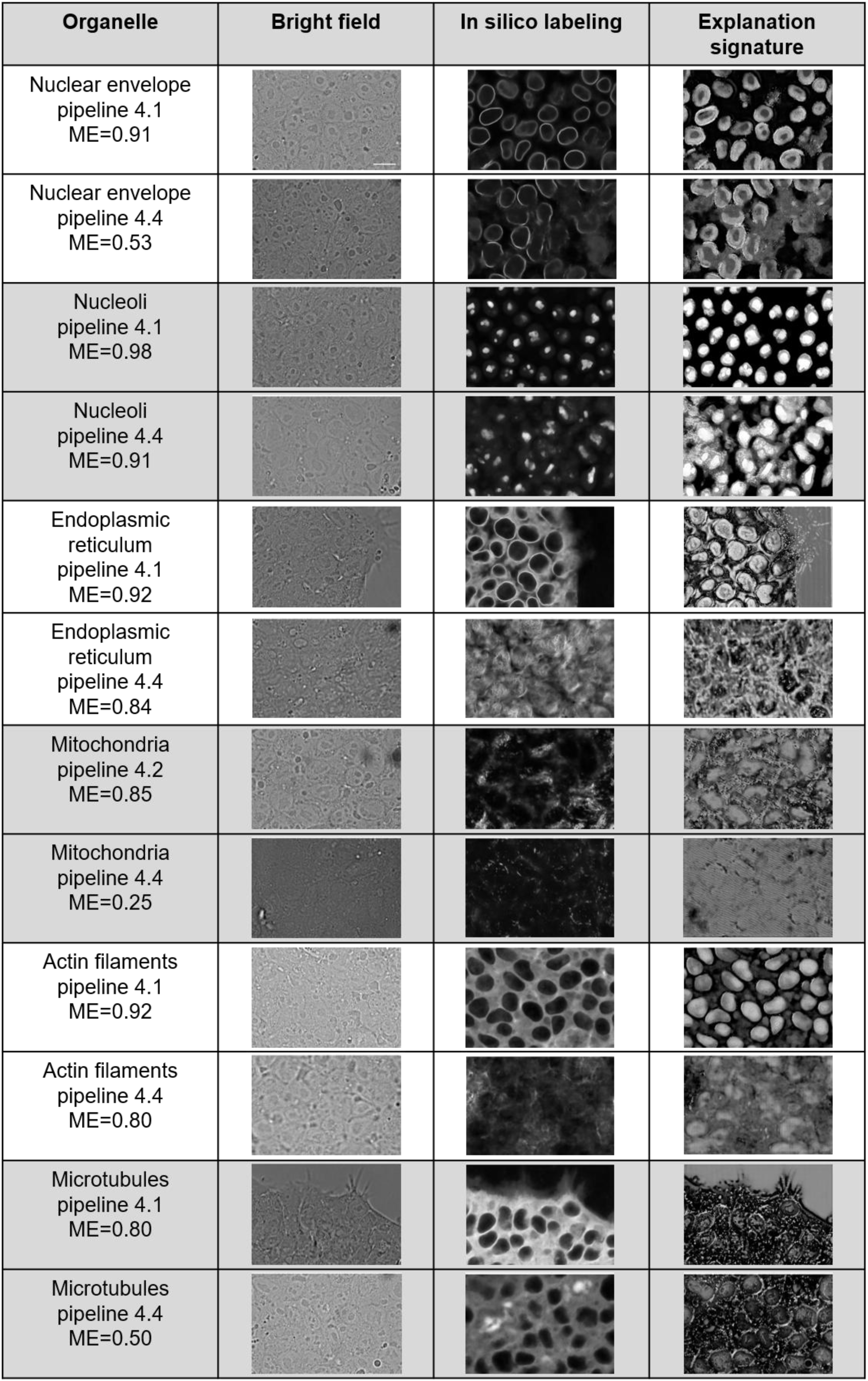
Visual assessment of potential batch effects in imaging pipeline 4.4 across organelles: in silico labeling predictions and their corresponding Mask Interpreter’s explanation signatures and mask efficacies. Each paired row shows two representative fields of view of a specific organelle and imaging pipelines (left-to-right): the bright-field image, the corresponding in silico labeling prediction, and the Mask Interpreter’s explanation signature. Consistent with the quantitative results in Fig. S8, for some organelles pipeline 4.4 shows lower Mask efficacies (ME) that were associated with visual degradation of the in silico labeling quality, and non-stereotypical explanation signatures. Scale bar = 15 μm.

**Figure S10.**
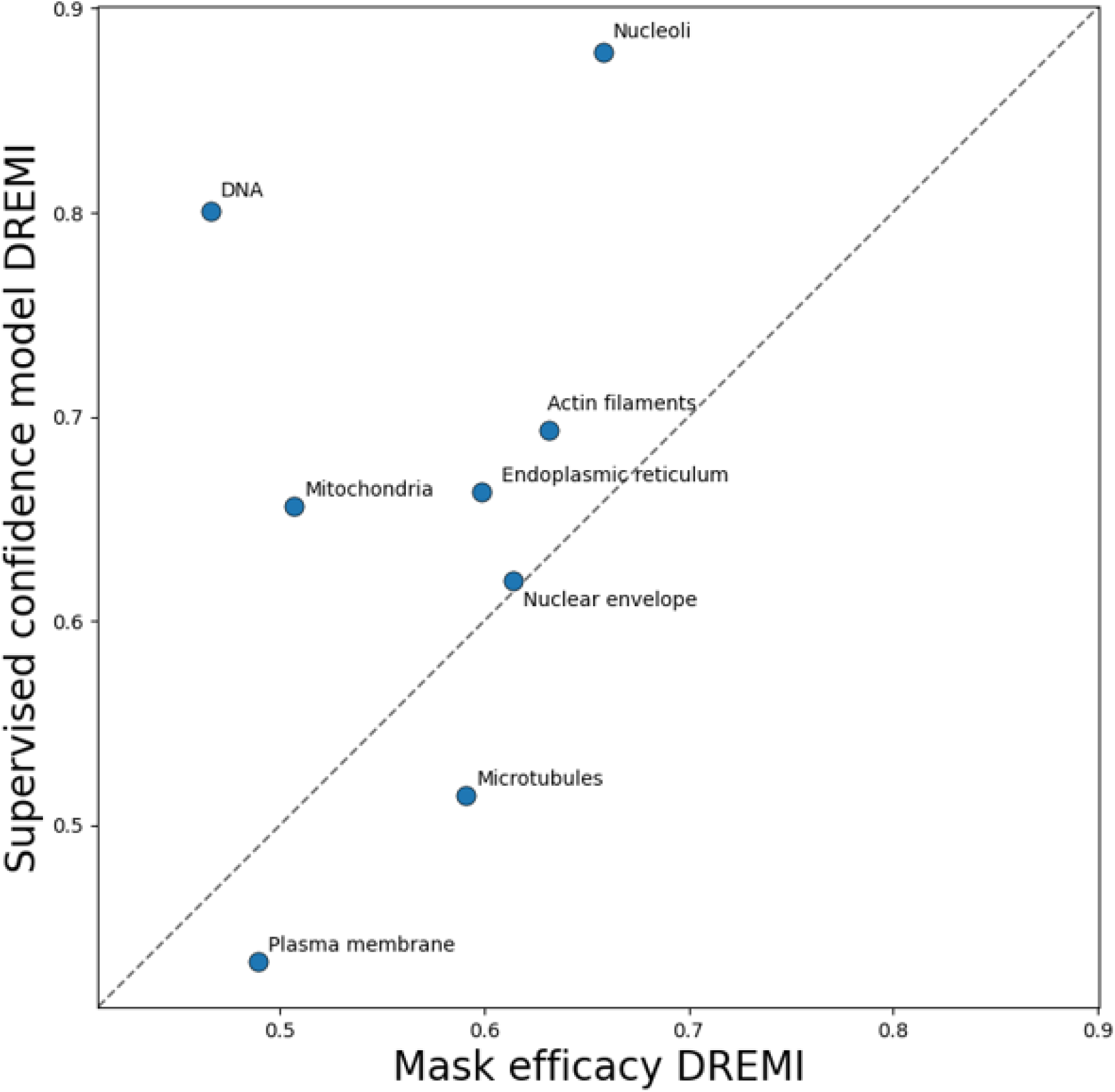
DREMI comparison between supervised confidence and mask efficacy. Each point represents one organelle and shows the DREMI between the true patch-level PCC and either mask efficacy (x-axis) or the supervised confidence model prediction (y-axis). The dashed line is the diagonal Y = X, indicating equal performance; points above the line favor the supervised model, whereas points below the line favor mask efficacy.

**Figure S11.**
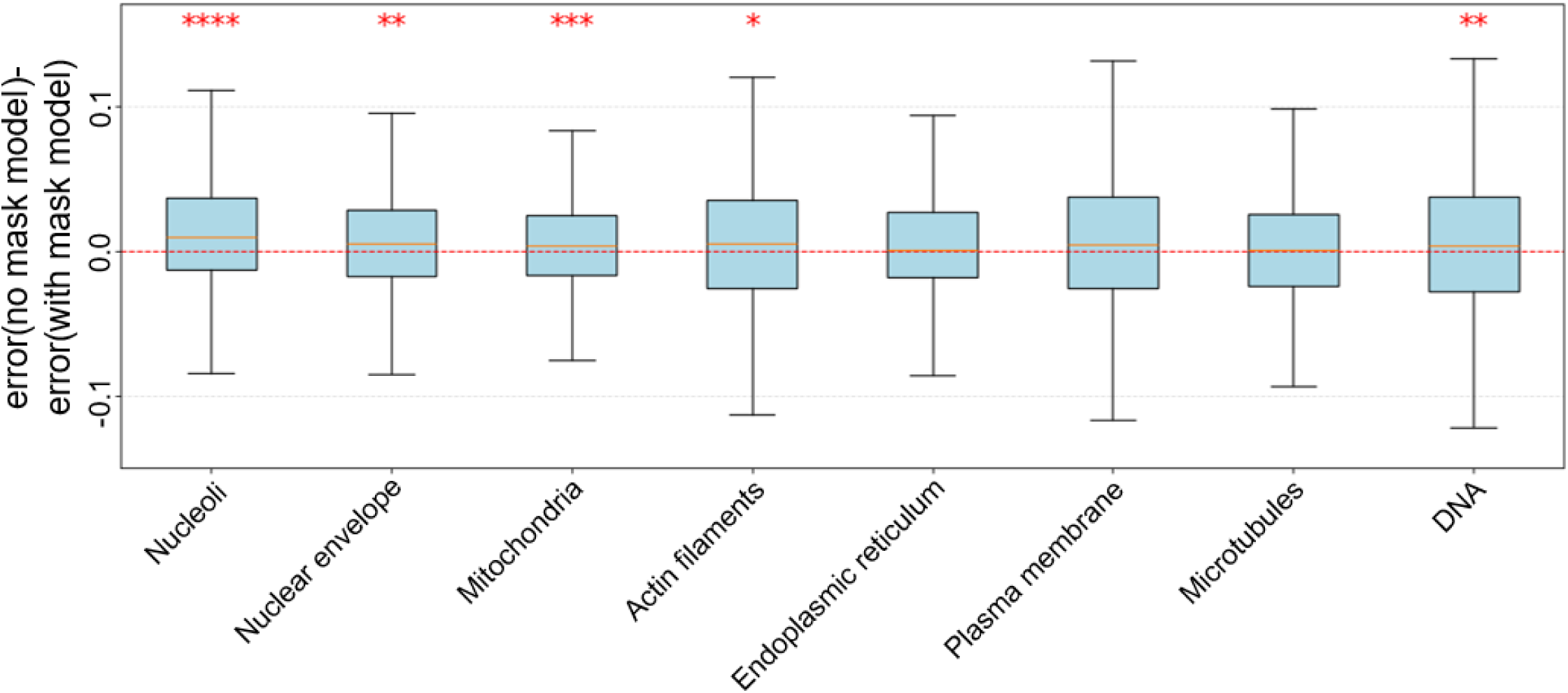
Ablation study comparing supervised confidence models with and without importance mask. For each organelle, performance differences were computed in a paired model-comparison analysis. The y-axis shows the difference in absolute PCC error between the compared confidence measures/models. Values above zero indicate better performance of the confidence model with the importance mask as input. The statistical test evaluates whether the observed performance difference between the compared confidence measures/models is larger than expected by random variation. *P < 0.05, **P < 0.01, ***P < 0.001, ****P < 0.0001 . Box plots show the median and interquartile range (IQR); whiskers extend to the most extreme values within 1.5 × IQR. The red dashed horizontal line marks zero difference between the compared confidence measures/models.

**Figure S12.**
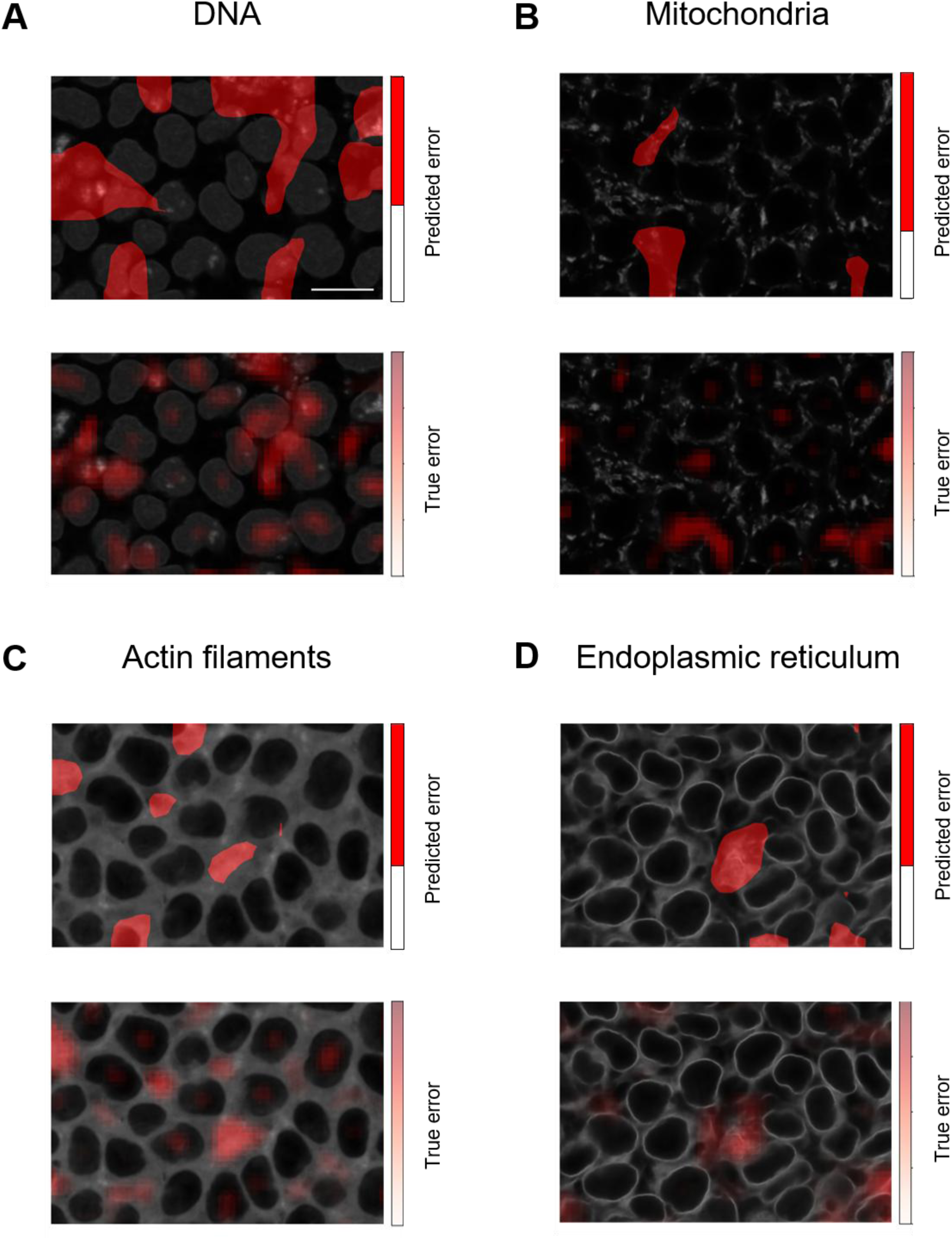
Supervised confidence modeling identifies image regions with poor in silico labeling at inference. Overlay visualizations showing representative in silico labeling confidence scores (top) and the corresponding ground truth in silico labeling errors (bottom) on top of the in silico labeling predictions for the DNA (A, threshold=0.43), mitochondria (B, threshold=0.3), Actin Filaments (C, threshold=0.37) and Endoplasmic Reticulum (D, threshold=0.37) confidence is measured by 1 minus the PCC between in silico prediction and ground truth fluorescence image. Red regions indicate low confidence (top) and poor in silico labeling performance (bottom). Visual assessment indicates association between low confidence and poor in silico labeling performance. Scale bar = 15 μm.

**Figure S13.**
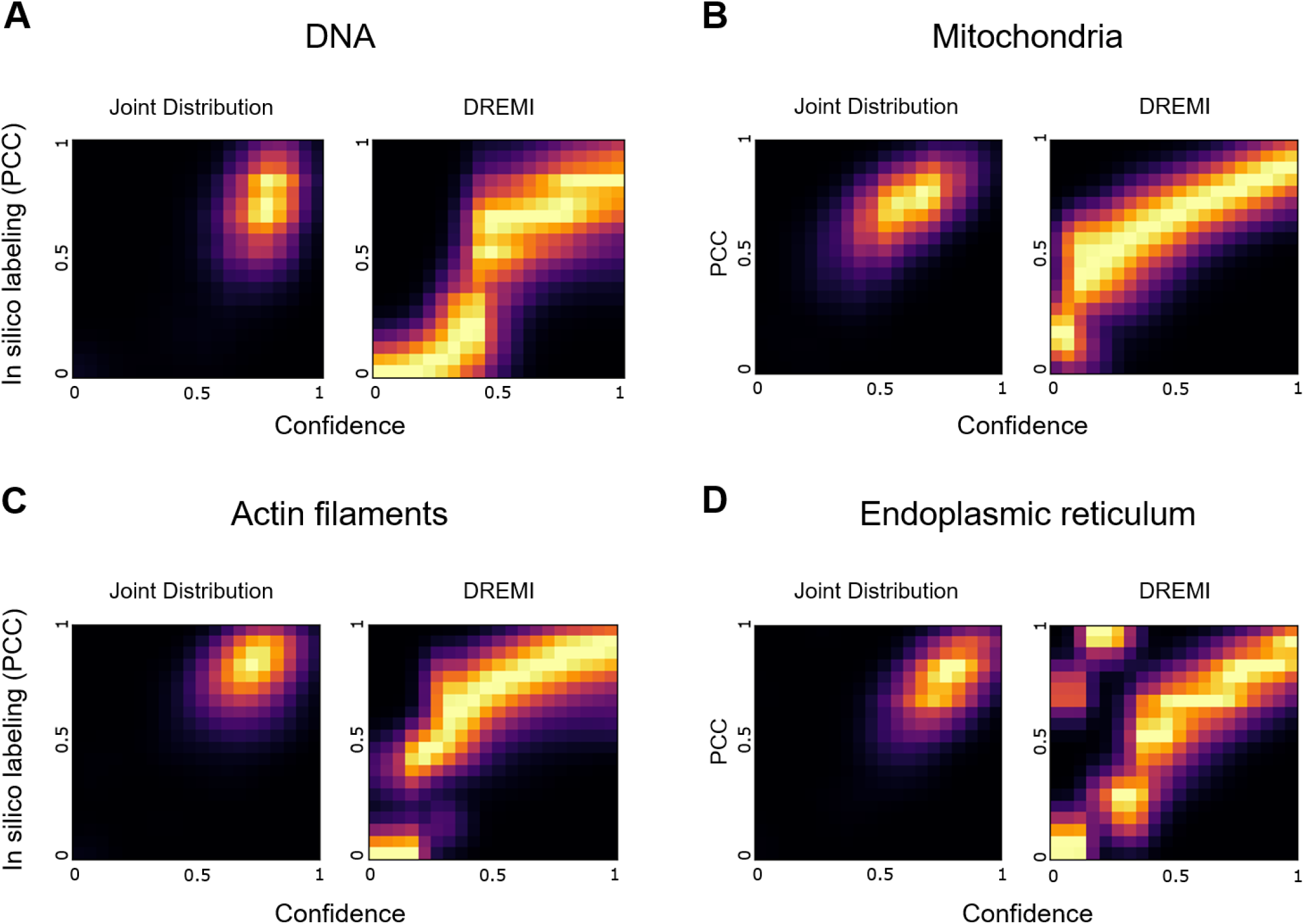
The confidence model accurately predicts in silico labeling quality across organelles. Systematic evaluation of the confidence model for the: DNA (A, 810 patches), mitochondria (B, 972 patches), actin filaments (C, 540 patches) and the endoplasmic reticulum (D, 378 patches) using DREMI analysis of the predicted versus ground truth in silico labeling PCC. DREMI scores (A to D): 0.80, 0.66, 0.69, 0.66.

**Figure S14.**
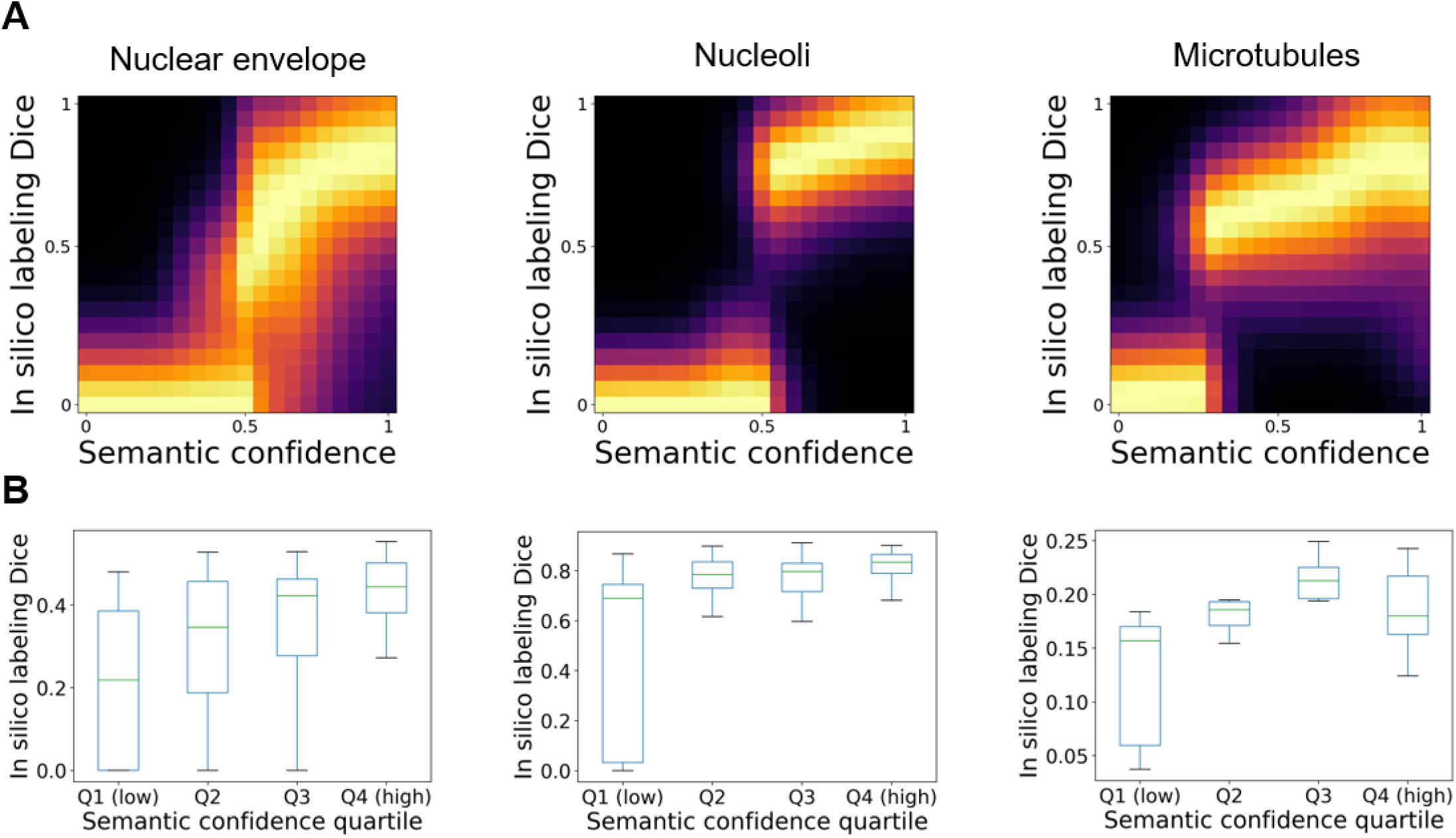
Single cell semantic confidence associates with the Dice Coefficient between the segmented in silico labeling and the corresponding segmented fluorescence image. Analysis presented for three organelles, in accordance with Fig. 7B-C: Nuclear Envelope (143 cells), nucleoli (176 cells), and microtubules (33 cells). (**A**) Conditional density (DREMI) between the single cell semantic confidence (x-axis) and the Dice Coefficient (left-to-right): 0.47, 0.80, 0.51. (**B**) Cells were grouped into quartiles (Q1–Q4) based on their semantic confidence scores, with Q1 corresponding to the lowest confidence and Q4 to the highest. Boxes indicate the interquartile range, center lines indicate the median, and whiskers extend to 1.5× the interquartile range (outliers not shown). Dice coefficient was higher in the pooled Q2–Q4 cells than in Q1 cells for all three organelles: Nuclear Envelope (median: 0.420 versus 0.219; one-sided Mann–Whitney U = 2897.5, p = 3.0 × 10⁻⁶; Cliff’s δ = 0.504), nucleoli (0.803 versus 0.690; U = 4822, p = 2.85 × 10⁻¹¹; Cliff’s δ = 0.660), and microtubules (0.193 versus 0.157; U = 182, p = 1.48 × 10⁻³; Cliff’s δ = 0.685).

**Figure S15.**
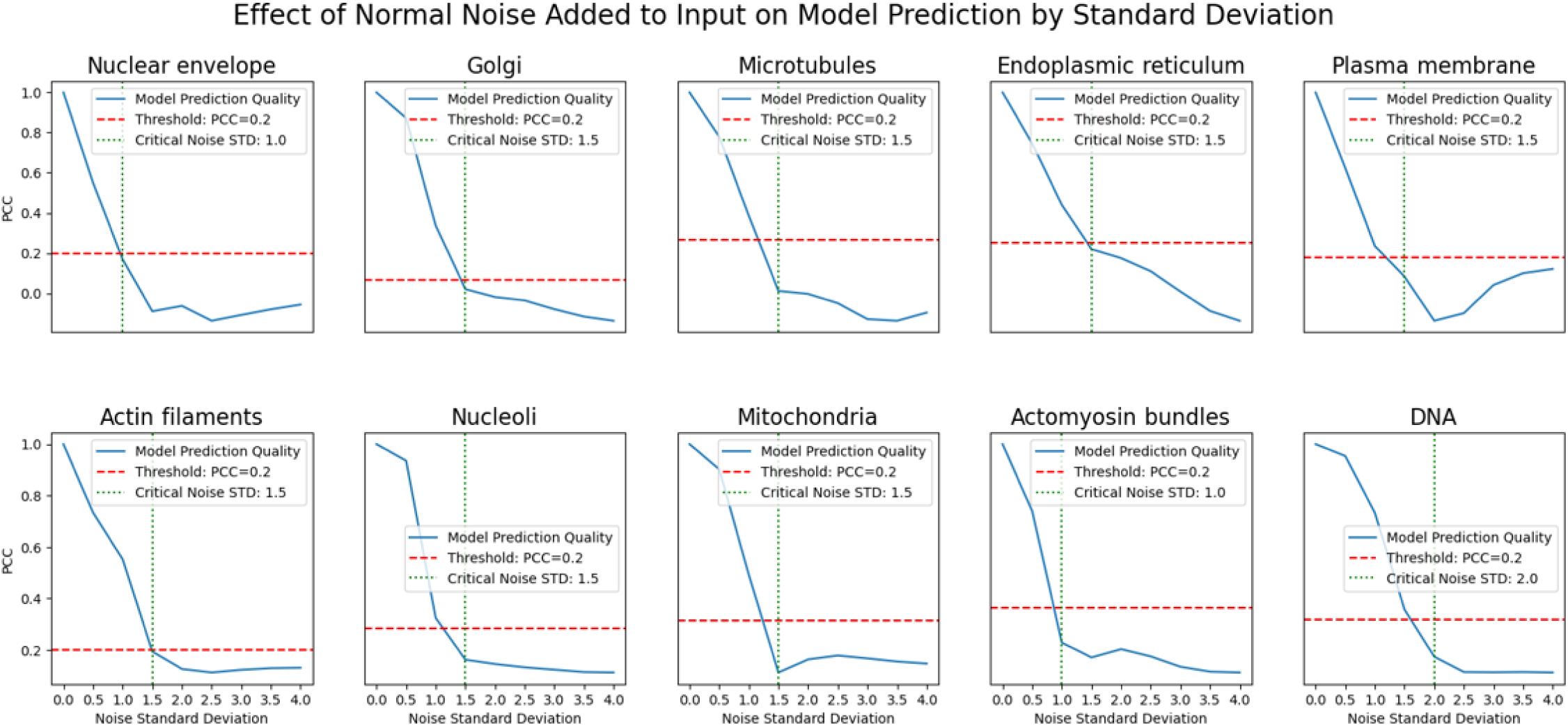
Determination of the Gaussian’s noise standard deviation (σ). Organelle-specific plots show the Pearson correlation coefficient (PCC) between in silico labeling predictions from the unperturbed and the noisy bright-field images as a function of the Gaussian noise (σ) included. For each organelle we selected σ (vertical green dashed lines) as the smallest value where the PCC drops below 0.2 (horizontal red dashed line), ensuring sufficient noise to identify non-essential regions without overwhelming the image.

**Figure S16.**
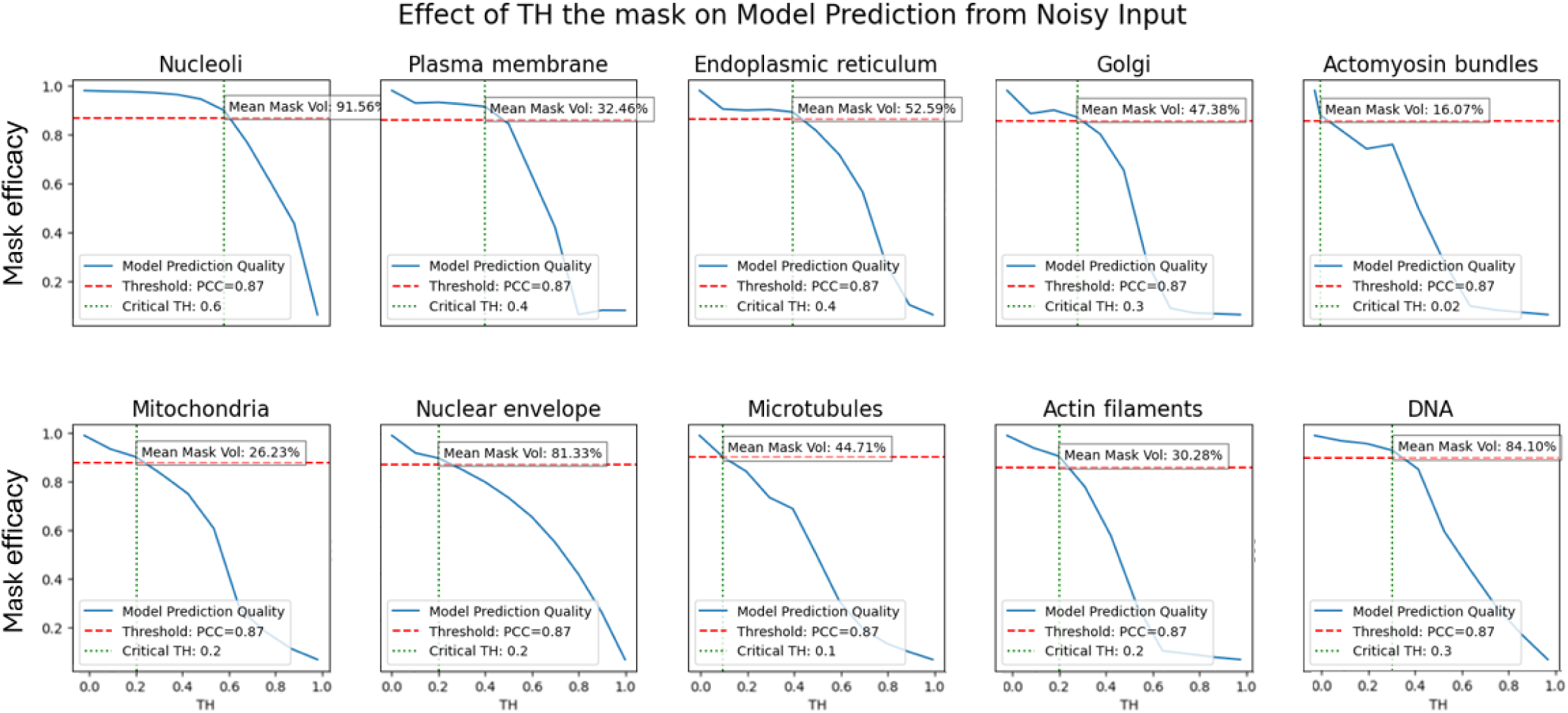
Threshold determination for generating binary explanation signatures. For each of the ten organelles examined, thresholds ranging from 0.0 to 1.0 were applied to the importance masks produced by the Mask Interpreter. Voxels below the threshold were considered non-essential and were noised. The mask efficacy between the in silico labeling from the noisy and unperturbed bright-field image was computed. The chosen threshold (vertical green dashed line) for each organelle corresponds to the first point at which the mask efficacy fell below 0.87 (horizontal red dashed line), ensuring that the resulting binary explanation signature minimizes coverage while maintaining prediction fidelity.

### Supplementary Tables

**Table S1.**
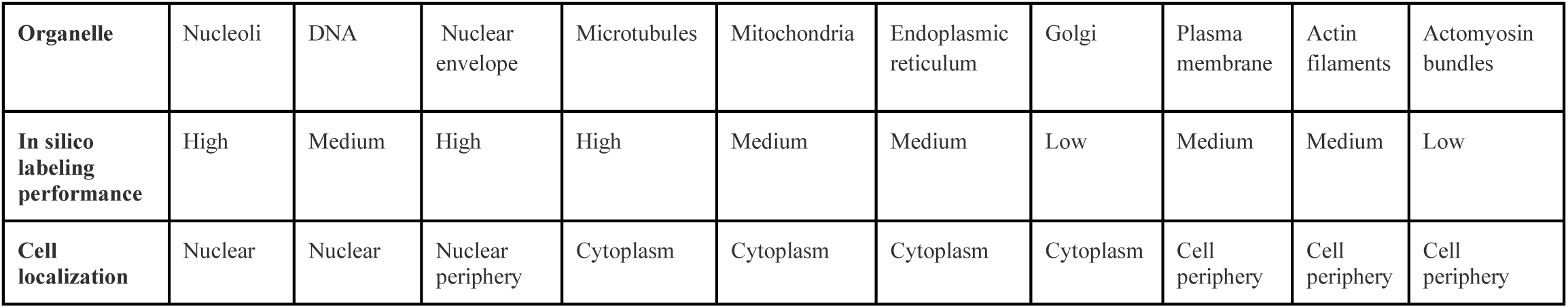
Summary of the organelles’ localization and their in silico labeling performance. Ten organelles were selected to span a range of structures, cellular localization^38^, and in silico labeling performances^9^.

| Organelle | Nucleoli | DNA | Nuclear envelope | Microtubules | Mitochondria | Endoplasmic reticulum | Golgi | Plasma membrane | Actin filaments | Actomyosin bundles |
| --- | --- | --- | --- | --- | --- | --- | --- | --- | --- | --- |
| In silico labeling performance | High | Medium | High | High | Medium | Medium | Low | Medium | Medium | Low |
| Cell localization | Nuclear | Nuclear | Nuclear periphery | Cytoplasm | Cytoplasm | Cytoplasm | Cytoplasm | Cell periphery | Cell periphery | Cell periphery |

**Table S2.** Interpretation of the Mask Interpreters’ explanation signatures. The table summarizes what were the regions in the bright-field image that drive the in silico labeling according to our interpretation of the corresponding explanation signatures.

| Organelle | Nucleoli | DNA | Nuclear envelope | Microtubules | Mitochondria | Endoplasmic reticulum | Golgi | Plasma membrane | Actin filaments | Actomyosin bundles |
| --- | --- | --- | --- | --- | --- | --- | --- | --- | --- | --- |
| Interpretation of the Mask Interpreter's explanation signature | Organell e and localized spatial context around it | Nuclei and broad spatial context surrounding them | Nucleus and spatial context around the organelle | Negative spatial context within large image regions that do not include the organelle | organelle along with the spatial context of most of the cytoplasm | Negative spatial context within large image regions that do not include the organelle | Negative spatial context within large image regions that do not include the organelle | Negative spatial context within large image regions that do not include the organelle | Negative spatial context within large image regions that do not include the organelle | Negative spatial context within large image regions that do not include the organelle |

**Table S3.** Benchmarking confidence models using the Mean squared error (MSE) metric. MSE between the predicted and the true in silico labeling quality for each organelle. Benchmarking mask efficacy, the supervised model that received the in silico labeled patch as input, and the supervised model that received both the in silico labeled patch and the Mask Interpreter-derived importance mask as input. Bold indicates the best-performing model for each organelle (lowest MSE).

| Organelle | Nucleoli | DNA | Nuclear envelope | Microtubules | Mitochondria | Endoplasmic reticulum | Plasma membrane | Actin filaments |
| --- | --- | --- | --- | --- | --- | --- | --- | --- |
| Mask efficacy | 0.1090 | 0.0622 | 0.0093 | 0.0189 | 0.0242 | 0.0339 | 0.0900 | 0.0203 |
| Confidence model (in silico labeling) | 0.0093 | 0.0146 | 0.0048 | 0.0062 | 0.0026 | 0.0049 | 0.0105 | 0.0089 |
| Confidence model (in silico labeling + importance mask) | **0.0061** | **0.0131** | **0.0043** | **0.0060** | **0.0024** | **0.0044** | **0.0100** | **0.0081** |

**Table S4.** Benchmarking confidence models using the Pearson correlation coefficient (PCC) metric. PCC between the predicted and the true in silico labeling quality for each organelle. Benchmarking mask efficacy, the supervised model that received the in silico labeled patch as input, and the supervised model that received both the in silico labeled patch and the Mask Interpreter-derived importance mask as input. Bold indicates the best-performing model for each organelle (highest PCC).

| Organelle | Nucleoli | DNA | Nuclear envelope | Microtubules | Mitochondria | Endoplasmic reticulum | Plasma membrane | Actin filaments |
| --- | --- | --- | --- | --- | --- | --- | --- | --- |
| Mask efficacy | 0.8720 | 0.2530 | 0.6230 | 0.6740 | 0.5660 | 0.6310 | 0.5020 | 0.6250 |
| Confidence model (in silico labeling) | 0.8690 | 0.7070 | 0.6660 | 0.7300 | 0.7100 | 0.7020 | 0.4490 | 0.6220 |
| Confidence model (in silico labeling + importance mask) | **0.9160** | **0.7360** | **0.7050** | **0.7310** | **0.7290** | **0.7150** | **0.5220** | **0.6540** |

**Table S5.** Benchmarking confidence models using the Coefficient of determination (R-squared) metric. R-squared between the predicted and the true in silico labeling quality for each organelle. Benchmarking mask efficacy, the supervised model that received the in silico labeled patch as input model, and the supervised model that received both the in silico labeled patch and the Mask Interpreter-derived importance mask as input. Bold indicates the best-performing model for each organelle (highest R-squared).

| Organelle | Nucleoli | DNA | Nuclear envelope | Microtubules | Mitochondria | Endoplasmic reticulum | Plasma membrane | Actin filaments |
| --- | --- | --- | --- | --- | --- | --- | --- | --- |
| Mask efficacy | 0.7076 | -1.1942 | -0.2023 | -0.5022 | -3.6639 | -2.9796 | -6.4430 | -0.5070 |
| Confidence model (in silico labeling) | 0.7504 | 0.4854 | 0.3834 | 0.5042 | 0.4939 | 0.4281 | 0.1345 | 0.3398 |
| Confidence model (in silico labeling + importance mask) | **0.8364** | **0.5382** | **0.4381** | **0.5225** | **0.5288** | **0.4774** | **0.1738** | **0.4003** |

